# Mite-induced migratory skin-like CD103^+^ ILC2s establish lung residency and impact immunity

**DOI:** 10.64898/2026.05.26.727651

**Authors:** Mi Lian, Tithi Roy, Lena Tama, Fanglue Peng, Konrad Knöpper, Longhui Qiu, Hong-Erh Liang, Georg Gasteiger, Roberto R. Ricardo-Gonzalez, Richard M. Locksley

**Affiliations:** Howard Hughes Medical Institute, University of California, San Francisco, CA, USA; Department of Medicine, University of California, San Francisco, CA, USA; Department of Dermatology, University of California, San Francisco, CA, USA; Department of Microbiology & Immunology, University of California, San Francisco, CA, USA; Wuerzburg Institute of Systems Immunology (WUESI), Max Planck Research Group at the Julius-Maximilians-University Wuerzburg, Wuerzburg, Germany; Biohub, San Francisco, CA, USA

## Abstract

Tissue-resident immune cells are critical for lung homeostasis and protection against inflammation. Here, we describe a novel population of migratory inflammatory CD103^+^ skin ILC2s (skILC2s) that accumulate in the lung vasculature following transient infestation with ubiquitous Demodex mites, hair follicle commensals that drive local skILC2 proliferation and entry into blood. With time, ex-skILC2s become tissue resident in the lung, occupy previously described immune cell adventitial niches and respond earlier to lung perturbations than endogenous lung ST2^+^ ILC2s to alter the trajectories of a subsequent immune response. Thus, like transiting ex-gut ILC2s described after small intestinal infection by helminths and protists, ex-skILC2s migrate post-birth in response to pathobionts to change the inflammatory milieu of the lung, revealing a common paradigm by which internal mucosal responses become shaped by ILC2s from barrier epithelia colonized post-birth.

**Highlights:**

- Demodex mites infestation drives local expansion of CD103^+^ skin ILC2s that egress and enter the circulation
- Migratory CD103^+^ skin-like ILC2s infiltrate into lung, adapt to the local environment and establish long-term residency
- Prior Demodex mite exposure re-organizes the lung ILC2 landscape
- The distinct cytokine and receptor repertoire of ex-skILC2 alters lung immunity

## Introduction

Tissue-resident immune cells are involved in basal homeostasis and in initiating responses to perturbations that upset homeostasis.^1, 2, 3, 4, 5, 6, 7^ Mechanisms that establish the normal composition, positioning, survival and activation of tissue-resident immune cells are critical for orchestrating the complex local and systemic cell circuits that promote homeostasis, protection, repair and memory.^8, 9, 10^ Developmentally, innate lymphoid cells (ILCs), including NK, lymph node tissue-inducer cells (LTi), and Group 1, 2 and 3 ILCs, constitute the earliest wave of tissue-resident lymphoid cells.^11, 12, 13^ While lacking the antigen-specific receptors of adaptive lymphocytes, Group 1-3 ILCs share transcriptional and cytokine effector programs with Th1, Th2 and Th17 cells, respectively, but differentiate in tissues, revealing shared core responses implicated in homeostasis and initiation of systemic immune responses to different types of inflammation.^14, 15, 16, 17^ Further understanding of mechanisms underlying adaptation of ILCs to tissue are likely to extend to processes by which adaptive T cells differentiate to become resident effector and memory cells in diverse tissues and under different conditions throughout life.

In small intestine, Group 2 innate lymphoid cells (ILC2s) constitutively produce cytokines IL-13 and IL-5 in response to the regulated release of alarmins, predominantly IL-25 but also TSLP, during homeostasis^18, 19, 20, 21^ but also following gut colonization by certain protists and helminths.^22, 23, 24^ Although homeostatic responses remain localized, parasite infestation drives ILC2 proliferation, entry into the blood and accumulation in the lung with impact on the subsequent lung immune response to heterologous challenges.^25, 26, 27^ Indeed, repeated doses of IL-25 can alone drive accumulation of memory gut-like ILC2s in lung that potentiate protection against later challenges.^20, 26, 27^ Thus, developmentally positioned gut ILC2s migrate upon colonization by pathobionts post-birth and affect immunity at the lung mucosa to control subsequent insults. Whether this is a unique property of the highly colonized bowel or whether this process is broadly extended to other colonized barrier sites is not known.

Previously, we demonstrated that control of Demodex, a commensal mite that infests mammalian hair follicles, requires IL-13 and the type II IL-4 receptor, IL13Rα1 and IL4Rα; absence of any of these resulted in uncontrolled mite replication and skin ILC2 (skILC2) expansion, acquisition of an inflammatory phenotype, and appearance in the blood.^28^ Here, we co-housed Demodex-infested type 2-immunodeficient mice with uninfected immunocompetent neonatal or adult mice for 2 or 3 weeks before separating the animals to allow mite clearance to assess the impact of transient hair follicle colonization on the ILC2 landscape of the lungs. As in helminth-infected mice, we show that Demodex colonization drives expansion of an inflammatory CD103^+^ skILC2 population that egresses into blood and establishes tissue residency in lung adventitial niches previously described to house resident immune cells.^29, 30, 31^ Further, diverse lung challenges revealed that ex-skILC2s have early activation kinetics and an altered cytokine repertoire that impact tissue, suggesting a broader paradigm by which ILC2s from external barriers ferry information to internal mucosa of the lung to enhance host resilience.

## Results

### CD103^+^ skin-like ILC2s accumulate in the lungs of Demodex-infested IL4Rα^-/-^ mice

At steady state, most ILC2s in specific-pathogen free (SPF) C57BL/6 mice are ST2^-^IL18R1^+^ CD103^+^ in skin (subcutaneous fat was removed) and ST2^+^IL18R1^-^ CD103^-^ in lung (**Figure 1a**).^32, 33^ We previously described expansion of ILC2s in skin and blood of IL13Rα1^-/-^, Il4/Il13^-/-^ and IL4Rα^-/-^ mice that are unable to control cutaneous Demodex.^28^ Additionally, we noted accumulation of lung ILC2s in mite-infested IL4Rα-deficient and IL-13-deficient mice with the ST2^-^IL18R1^+^CD103^+^ skin-like phenotype (**Figure 1b, S1a, b**). Pharmacologic clearance of mites from infested Il4/Il13^-/-^ and IL4Rα^-/-^ mice resulted in significant contraction of ILC2s in skin, but persistence of CD103^+^ skin-like ILC2s in lungs (**Figure S1c, and data not shown**).

**Figure 1.**
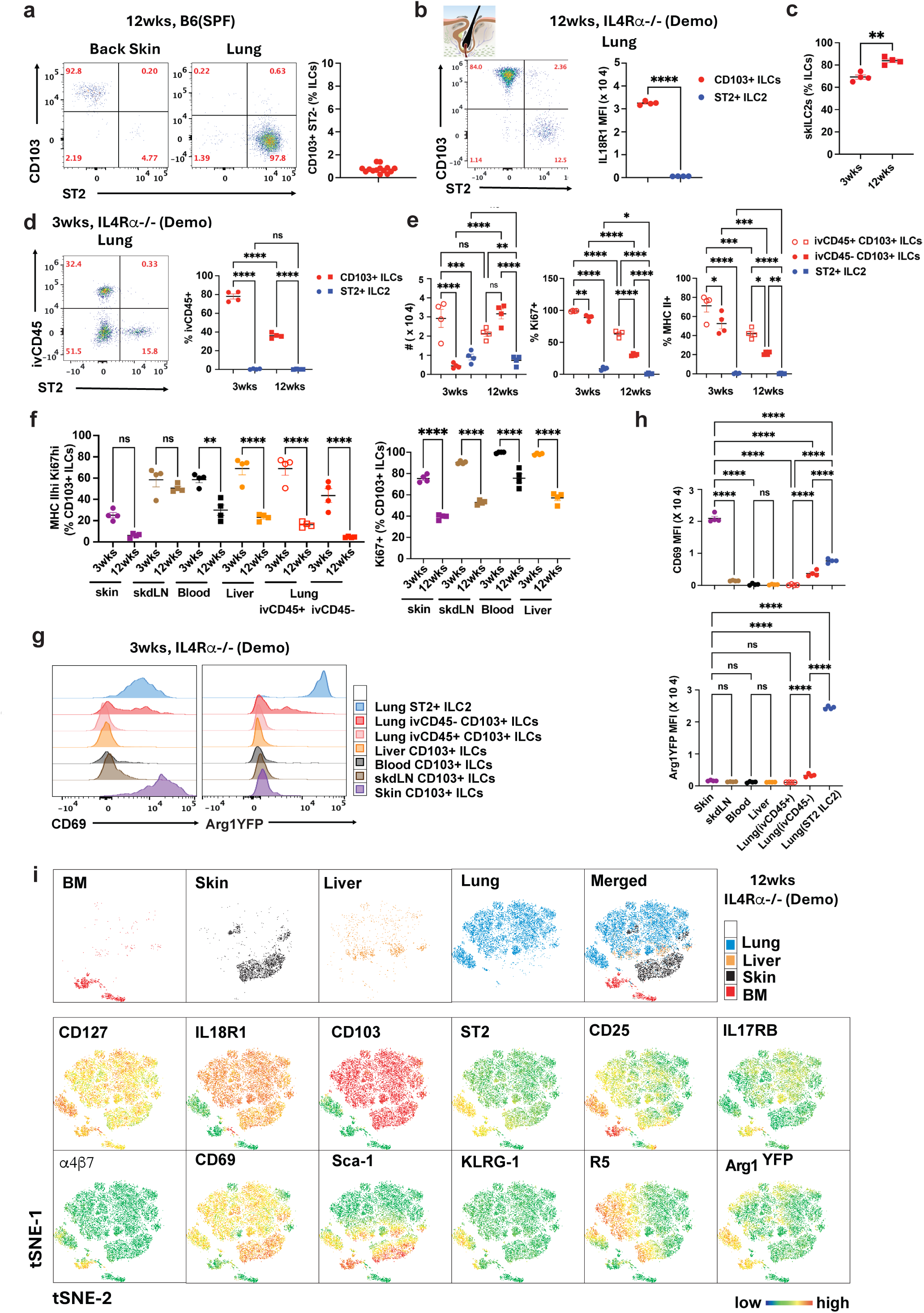
Skin-like CD103^+^ ILC2s accumulate in lungs of Demodex-infected IL4Rα^-/-^ mice. **a.** Expression of CD103 and ST2 on back skin (subcutaneous fat removed) and lung CD127^+^ CD90.2^+^ ILCs in SPF B6 mice. **b.** FACS plot of lungs from Demodex, hair follicle mite-infected IL4Rα^-/-^ mice (Demo) at age of 12 weeks (12wks) showing skin-like ST2^-^ CD103^+^ ILCs (left), and graph for the expression of IL18R1 on CD103^+^ ILCs and ST2^+^ ILC2 (right). **c.** Percentage of skin-like CD103^+^ ILCs in IL4Rα^-/-^ (Demo) lungs at age of 3wks and 12wks. **d.** ercentage of ivCD45-labelled CD103^+^ ILCs and lung ST2^+^ ILC2 in IL4Rα^-/-^ (Demo) lungs at age of 3wks. **e.** Cell number, percentage of Ki67^+^ and MHC II^+^ expression on ivCD45^+^ CD103^+^ ILCs, ivCD45^-^ CD103^+^ ILCs, ST2^+^ ILC2 in IL4Rα^-/-^ (Demo) lungs at age of 3wks and 12wks. **f.** Percentage of MHC II^hi^ Ki67^hi^ and Ki67^+^ of CD103^+^ ILCs in tissues from IL4Rα^-/-^ (Demo) mice. **g, h.** Histogram (**g**) and graph (**h**, mean fluorescence intensity, MFI) comparison of CD69 and Arg1YFP expression on Il5^RFP+^ CD103^+^ ILCs among indicated tissues compared with ivCD45^+^ vs ivCD45^-^ CD103^+^ ILCs and ST2^+^ ILC2 in lungs of IL4Rα^-/-^ (Demo) mice at age of 3wks. **i.** tSNE plots generated from spectral flow cytometry for ILCs from lung, liver, bone marrow (BM), and ear skin from IL4Rα^-/-^ (Demo) mice and expression of the indicated markers for the merged tissue-specific ILCs. *p< 0.05, **p< 0.01, ***p<0.001, ****p<0.0001 by unpaired t-test **(c,d)** or one-way ANOVA **(h, i)**. n=4 biological replicates/group, representative of 2-3 independent experiments.

By 3 weeks (wks) of age, offspring of mite-infested IL4Rα^-/-^ mice accumulated skin-like ILC2s in lung (**Figure 1c**) and 80% were labelled by intravenous anti-CD45 antibody (**Figure 1d),** suggesting enrichment at vascular niches distinct from resident parenchymal ivCD45^-^CD103^-^IL18R1^-^ST2^+^ lung ILC2s. Over time, the percentage of ivCD45^+^ skin-like ILC2s decreased (**Figure 1d**) and parenchymal ivCD45^−^ CD103^+^ ILC2s increased coincident with a decrease in markers for proliferation (Ki67^+^) and activation (MHC II^+^)^33, 34, 35^ (**Figure 1e**), and consistent with adaptation to lung residency.^36, 37^ Further exploration of skin-like CD103^+^ ILC2 redistribution in these mice revealed CD103^+^ ILC2s present in skin-draining lymph nodes (skdLN), blood, liver, lung and bone marrow (BM), but few in small intestine lamina propria (siLP) (**Figure 1f, i, S1d-f**). At 3 wks of age, most CD103^+^ ILC2s in peripheral tissues displayed an activated MHC II^hi^ Ki67^hi^ phenotype and did not express the residency marker CD69^38, 39^ or tissue-imprinted arginase-1 (**Figure 1g**); the latter, while not expressed by resident skILC2s, is constitutive in other resident ILC2 populations including in the lung.^32, 40^ Whereas CD103^+^ ILC2s in peripheral non-skin sites remained CD69^−^ Arg1^YFP^-negative, lung ivCD45^−^ CD103^+^ ILC2s began to express CD69 and Arg1^YFP^ (**Figure 1g, h**). We used multiparameter spectral cytometry to compare the phenotypes of CD103^+^ skin-like ILC2s across non-skin sites, ST2^+^ lung resident ILC2s and CD103^+^ local skin ILC2s from Demodex-mite infested 12-wk-old IL4Rα^-/-^ mice (**Figure 1i**). Tissue-distributed CD103^+^ skin-like ILC2s expressed IL18R1, low levels of CD25, IL17RB, and CD69 and did not express KLRG-1 or the gut-homing integrin α4β7.^41^ Bone marrow ILCPs, ILC2Ps and immature ILC2s remained immature and did not express activation markers such as KLRG-1, CD103 and MHC II^33^ (**Figure 1i, and data not shown**), or IL-5, as assessed by expression of IL5^RFP^.^32, 42, 43^

### Neonatal exposure to mites drives lung adaptation by a persisting population of skin-like ILC2s

Because IL4Rα^+/-^ mice spontaneously clear Demodex,^28^ we bred Il5^RFP^ (R5) or WT C57BL/6 mice with infected IL4Rα^-/-^Arg1^YFP^ mice and moved the weaned, heterozygote offspring to clean cages after 21 days to assess effects of self-limited mite infestation on lung ILC2s (**Figure 2a**). Although unexposed WT mice have very few CD103^+^ ILC2s in lungs, neonatal exposure led to accumulation of CD103^+^ ST2^-^ skin-like ILC2s that persisted at 6 wks of age (**Figure 2b**), which was similar to that in infected IL4Rα^-/-^ mice (**Figure 1b**). Thus, Demodex mite exposure during the neonatal period leads to accumulation of skin-like ILC2s in the lung even in the presence of IL4Rα sufficient to promote mite clearance. Over time, we noted the significant accumulation of ST2^+^CD103^+^ ILC2s in IL4Rα^+/–^ lungs (**Figure 2b, S2b, S2d)** with decreased IL18R1 and increased Arg1^YFP^ compared with ST2^−^CD103^+^ skin ILC2s (**Figure 2c**) and limited KLRG-1 and PD-1 as compared to lung resident ST2^+^CD103^−^ ILC2s (**Figure S2a)**. The percentage of ST2^+^CD103^+^ ILC2s increased inversely to the proportion of ivCD45^+^ CD103^+^ ILC2s (**Figure 2d, S2c**), such that the majority of ST2^+^CD103^+^ ILC2s lost ivCD45 labelling by 12 wks (**Figure S2c**) and began to express residency markers like CD69 and Arg1^YFP^ with reduced cell cycling (Ki67) (**Figure 2e, S2c-d)**. Lung CD103^+^ ILC2s produced a mixed type 2/3 cytokine profile including IL-5, IL-13 and IL-17A as compared to endogenous lung CD103^−^IL18R1^−^ST2^+^ ILC2s (**Figure 2f**), which produced IL-5, IL-13 and GM-CSF; both populations produced IL-2 and neither produced IL-22 as occurs among CD103^+^ ILC2s cells in the absence of IL4Rα **(Figure S2e).**^28, 33, 77^

**Figure 2.**
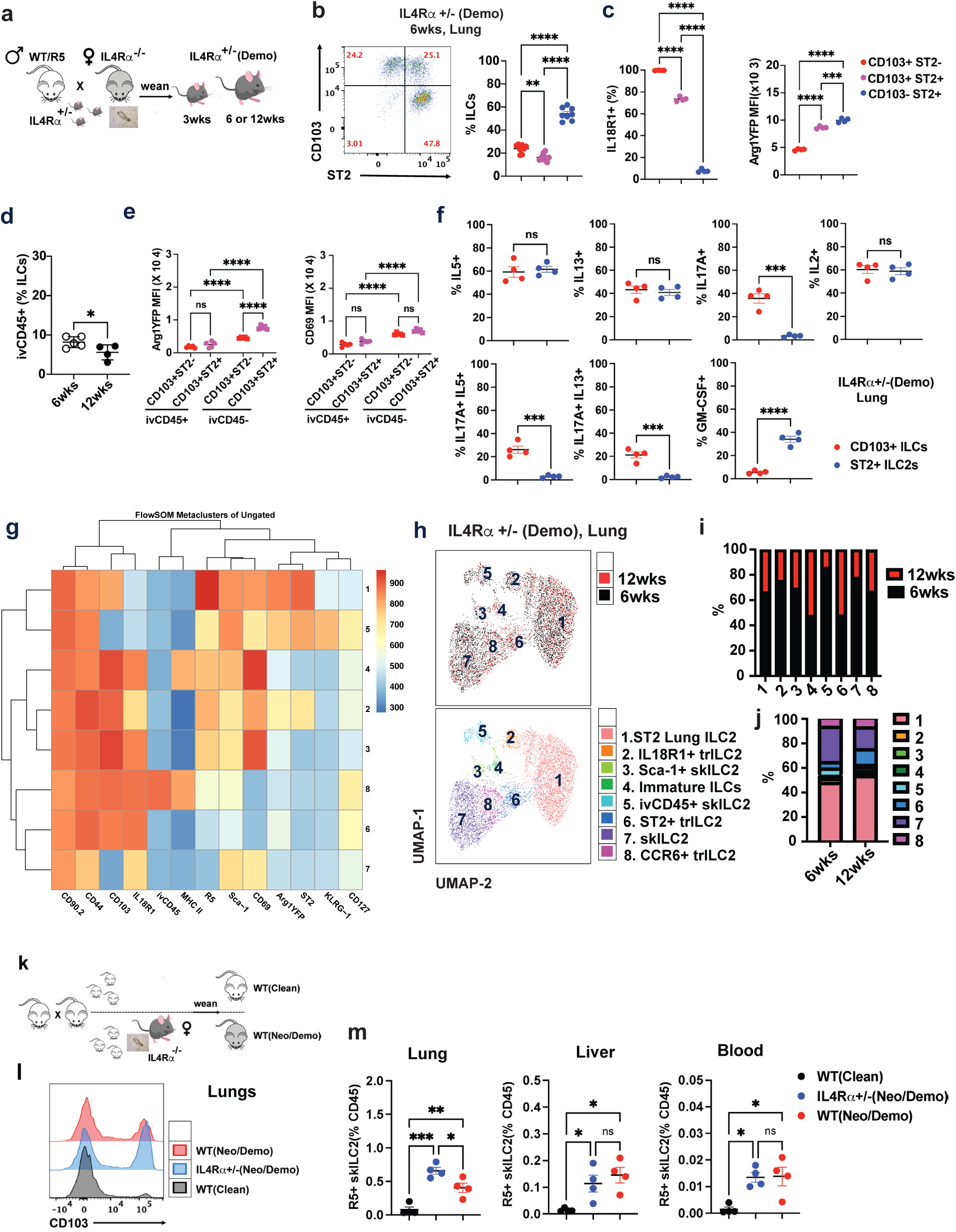
Neonatal exposure to Demodex mites leads to infiltration and adaptation of CD103^+^ skin-like ILCs in adult lungs. **a.** Breeding strategy to generate IL4Rα^+/-^ pups that were exposed to Demodex mite during the neonatal period, termed IL4Rα^+/–^ (Demo)**. b.** FACS plot and graph of percentage of CD127^+^ ILC subsets from IL4Rα^+/-^ (Demo) lungs, divided as CD103^+^ ST2^−^, CD103^+^ ST2^+^, and CD103^−^ ST2^+^ at 6wks of age. **c.** IL18R1 expression (left) and Arg1^YFP^ MFI (right) of indicated ILC subsets as in (**b**). **d.** Percentage of ivCD45^+^ ILCs in lungs of IL4Rα^+/–^ (Demo) mice at age of 6wks and 12wks**. e.** Arg1^YFP^ and CD69 MFI in indicated ILC subsets of IL4Rα^+/–^ (Demo) lungs at 6wks. **f.** Cytokine expression profiling of CD103^+^ ST2^-^ ILCs versus ST2^+^ CD103^-^ ILC2s from IL4Rα^+/–^ (Demo) lung upon *in vitro* PMA/Ionomycin re-stimulation. **g.** Heatmap of identified clusters (population, pop) by flowSOM analysis for merged spectral flow cytometry data from IL4Rα^+/–^ (Demo) lung ILCs. **h.** UMAP for merged samples (top) and indicated ILC subsets clusters (bottom) as in (**g**). **i, j.** Individual cluster (pop) comparison **(i)** and clusters (pop) distribution comparison **(j)** for ILCs from 6wks and 12wks IL4Rα^+/–^(Demo) lungs. **k.** Breeding strategy to generate wildtype (WT) pups with or without neonatal stage exposure to Demodex mites by co-housing with a Demodex-infected IL4Rα^-/-^ mouse, and separated from the mite donor after weaning, termed WT (Neo/Demo) or WT (Clean), respectively**. l.** Histogram for CD103 expression on ILCs in lungs from WT (Clean), WT(Neo/Demo) and IL4Rα^+/–^ (Neo/Demo) mice. **m.** Percentage of Il5^RFP+^ skILCs in lung, liver and blood of groups as in **(l).** *p< 0.05, **p< 0.01, ***p<0.001, ****p<0.0001 by unpaired t-test (**d, f**) or one-way ANOVA (**b, c, e, m**). n=4 biological replicates/group, representative of 2-3 independent experiments.

We used flowSOM analysis to characterize potential trajectories of lung ILC2s in neonatally mite-exposed IL4Rα^+/–^ mice at 6 and 12 wks of age (**Figure 2g-j, S2f-g)**.^44, 45^ As suggested by the earlier analysis, ivCD45^+^CD103^+^ lung ILC2s decreased inversely with appearance of CD103^+^ST2^-^ and then CD103^+^ST2^+^ populations that gradually lose CCR6,^32^ while gaining residency markers like CD69 (**Figure 2g-j, S2f-g)**. To confirm that these observations extend to mice with homozygous functional IL4Rα alleles, we co-housed Il5^RFP^Arg1^YFP^ mice with mite-infected IL4Rα^-/-^ mice for 3 wks during the neonatal period before separation (**Figure 2k**) and replicated the appearance of CD103-expressing skin-like ILC2s (**Figure 2I**) in lungs with increased expression of Arg1^YFP^ **(Figure S2h)**. Consistently, R5^+^ CD103^+^ skin-like ILC2s were also present in liver and blood (**Figure 2m**), suggesting that neonatal mite exposure of WT mice drives egress and systemic distribution of skin-like ILC2s, which can persist and infiltrate lung tissues to establish long-term residency.

### Demodex mite exposure alters the lung ILC2 landscape in adult mice

The neonatal period is associated with massive ILC2 expansion and tissue seeding by ILC2Ps, precursor populations that may have adaptive properties not shared by mature ILC2s.^33, 46, 47^ To assess whether Demodex can drive skin ILC2 systemic egress, migration and lung adaptation in adult mice, we co-housed adult C57BL/6 mice with Demodex-infected IL4Rα^-/-^ mice for 2 wks (designated 2C – 2 wks Co-housed) before separation and analysis after 4 and 8 wks (designated 2C4R and 2C8R, respectively – 4 or 8 wks), by which time mites are cleared (**Figure 3a**).^28^ As previously, we noted increased numbers of CD103^+^ST2^−^ skILC2s in lungs from 2C, 2C4R and 2C8R mice as compared to SPF-maintained control animals (**Figure 3b**). Further, the percentage of CD69^+^ skin-like ILC2s in the lung increased in 2C4R and 2C8R mice as compared with 2C mice suggesting their infiltration and retention as skin inflammation resolved (**Figure 3b**). In keeping with initial inflammation-driven migration, 2C mice demonstrated skin expansion of CD103^+^ skin ILC2s (**Figure 3e**) and significantly increased skin-like R5^+^ ILC2s in lungs (**Figure 3c, d**); 40% of these remained ivCD45^+^ and, among the ivCD45^−^ population, showed higher level of Arg1^YFP^ and CD69, but lower level of MHC II expression, consistent with adaptation to lung residency (**Figure 3d, S3a**). In the lung of 2C mice, approximately 60% of ILC2s were cycling and expressed MHC II and CXCR6;^48, 49^ liver CD103^+^ ILC2s were also Ki67^+^CD69^−^, and consistent with vascular distribution of inflammatory CD103^+^ skILC2s during the period of acute mite infestation (**Figure S3b, c**).

**Figure 3.**
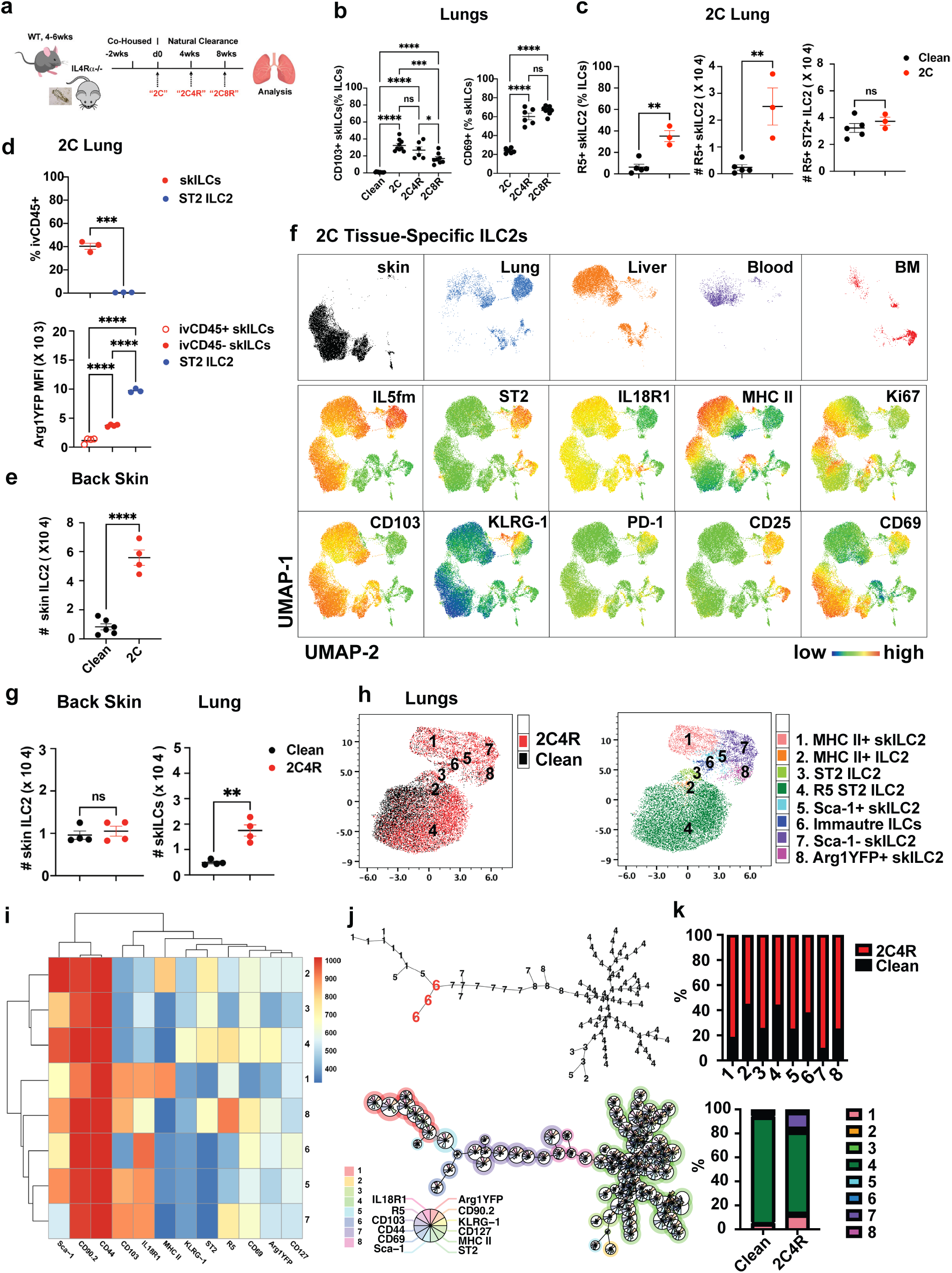
Demodex infection alters the lung ILC landscape in WT mice. **a.** Co-housing model to expose adult WT mice to Demodex-infected IL4Rα^-/-^ mice for 2wks, denoted as “2C”. After 2 weeks of exposure, WT mice are separated from the IL4Rα^-/-^ Demodex-infected mouse to allow naturally mite clearance for 4wks to 8wks, termed “2C4R” and “2C8R”, respectively. **b.** Percentage of skILC2s (left) and CD69^+^ skILC2s (right) in lungs from the indicated groups. **c.** Percentage of IL5^RFP+^ (R5) skILC2s and cell counts for CD103^+^ skILC2s and ST2^+^ ILC2 in 2C lungs. **d.** Percentage of ivCD45-labelled skILC2 and ST2^+^ ILC2, Arg1YFP MFI for indicated ILCs in 2C lungs. **e.** Cell counts for ST2^-^ CD103^+^ ILC2s (skILC2) in clean and 2C back skin. **f.** Sample distribution and expression for the indicated markers in UMAP for merged ILCs from indicated tissues of 2C R5Ai14 mice. **g.** Cell counts for skILC2s in back skin (subcutaneous fat removed) and skin-like ILCs in lungs of 2C4R mice. **h.** UMAP of merged ILC populations from clean and 2C4R, the indicated ILC clusters identified by flowSOM analysis. **i.** Heatmap of identified clusters (pop) by flowSOM analysis for merged lung ILCs in **(h). j.** Lineage trees generated by flowSOM analysis of samples in **(i),** represented as numbered populations (top) and by expression of markers for individual populations (bottom). **k.** Individual cluster (pop) comparison (**j**) and cluster distribution as a percentage of total (**k**) for ILCs from clean and 2C4R lungs. *p< 0.05, **p< 0.01, ***p<0.001, ****p<0.0001 by unpaired t-test (**c**, **d** (top), **e, g**) or one-way ANOVA (**b**, **d** (bottom)), n=4-5 biological replicates/group, representative of 2-3 independent experiments.

To assess further ILC2s in the acute phase of Demodex mite-driven egress, we compared ILC2s in skin with ILC2s from lung, liver, blood and bone marrow (BM) after 2 wks of mite exposure (**Figure 3f, S3d, e**). As expected, most IL5-fate mapped (IL5^fm+^) ILC2s in skin expressed CD69, whereas most non-skin CD103^+^ ILC2s exhibited an inflammatory Ki67^hi^MHCII^hi^CD69^lo^CD25^lo^ phenotype (**Figure 3f, S3d)**. Tissue ILC2s clustered into distinct populations including BM enriched for immature ILCs/ILCPs (CD103^−^ ST2^−^ IL18R1^+^) and immature ILC2s/ILC2Ps (ST2^+^ IL5^fm−^), whereas CD69^lo^MHCII^hi^ skin-like CD103^+^ ILC2s were found across peripheral tissues **(Figure S3d, e)**. Immature ILCs and ILC2Ps (**Figure S3d-g,** population (pop) 8) were also spread across tissues (**Figure S3f**), consistent with the concept that inflammation promotes circulation of less mature precursors.^33, 50, 51^ Using flowSOM to reveal potential lineage relationships among ILC subsets **(Figure S3h-j)** suggested that immature ILCs give rise to IL5^fm–^ST2^+^CD25^+^ ILC2Ps (pop 3), which further mature to lung resident IL5^fm+^ ILC2s (pop 2); activation drives further differentiation to a KLRG-1^hi^ inflammatory state (pop 4). Skin ILC2s (pop 1) can arise from immature ILCs (pop 8) and PD-1^hi^ ILC2Ps (pop 7) but can also differentiate into an inflammatory state (pop 5,6) with the potential to later achieve quiescence and gain ST2 expression, suggesting the capacity for lung adaptation and persistence.

After separation and resolution of mite infestation, the number of CD103^+^ skin ILC2s returned to normal in skin by 4 wks (2C4R) but remained significantly increased in the lung (**Figure 3g**). To compare the heterogeneity of lung ILC2s in non-mite-infested versus 2C4R mice, we merged spectral cytometry data (**Figure 3h, S3k**). Persisting lung CD103^+^ ILC2s with distinct cytokine/cytokine receptor profiles consistent with skin origin included subpopulations retaining skin-like (CD103^+^IL18R1^+^Arg1^YFP–^ST2^-^) or gaining lung-like (Arg1^YFP^ST2^+^) features (**Figure 3h-k, S3k**). To assess gene signatures among potential lung-adapting skin-like ILC2s, we reanalyzed our published single-cell RNA-seq data from skin and lung of IL5^RFP^ mice (**GSE117568**) (**Figure 4a**). Among 8 major clusters, lung cluster 6 expressed *Il18r1* and *Itgae*, as expressed by skin resident ILC2s (clusters 1,4,7), but also low levels of *Il1rl1* and *Arg1*, when compared to resident lung ILC2s (cluster 0,2,3,5) (**Figure 4a-d**). Lung ILC2s in cluster 6 also expressed transcripts for genes associated with migration (*S1pr1, S1pr4, Vim, Klf2, Cxcr4, Cxcr6*) and tissue infiltration (*Itgb1, Itgb2, Itgb7* but absent *Itga4* necessary for gut homing) (**Figure 4d, e**). As compared to other clusters of lung ILC2s, skin ILC2s differentially expressed genes (**Table S3**) and a core transcriptional network (*Itgae, Il18r1, Ccr6, Ccr8, Il13, Il17a, Cxcr4, S1pr4, H2-Ab1, Klf2, Tcf7* and *Itgb2*) that defined lung cluster 6 as a likely candidate for skin origin (**Figure 4f, g**). Of note, skILC2s expressed migratory-associated genes more highly than resident lung ILC2s at steady state (**Figure 4d, e**), consistent with low-level homeostatic movement of skin ILC2s to the lung with the potential to establish persistent residency, and consistent with prior studies in parabiotic animals ^36,47^ and with the homeostatic movement described among small intestine ILC2s.^18, 25, 26, 52^ Similar evidence was gained by reanalysis of data from unexposed and mite-exposed C57BL/6 mice (**GSE197983**) and corroborated gain of effector function (*Il13, Areg*), alterations in chemotactic capacity (*Ccr6, Ccr8, Cxcr4, Cxcr6*) and loss of *CD69* in the exposed group (**Figure 4i-k**). Taken together, these studies support a process for mite-induced differentiation of homeostatic skILC2s to inflammatory MHCII^+^ skILC2s with greater effector function and mobility that enter blood, become entrapped in the lung vasculature (ivCD45^+^) and, upon resolution of infection, move into lung tissue (ivCD45^−^) and begin adaptation by expression of Arg1 and ST2. Of note, the latter transition was more robust during neonatal as compared to adult infestation, in which a greater proportion of adapted cells derive from immature precursors from the lung and BM (**Figure 2g-j, S2f**), and in agreement with prior studies.^33, 51^

**Figure 4.**
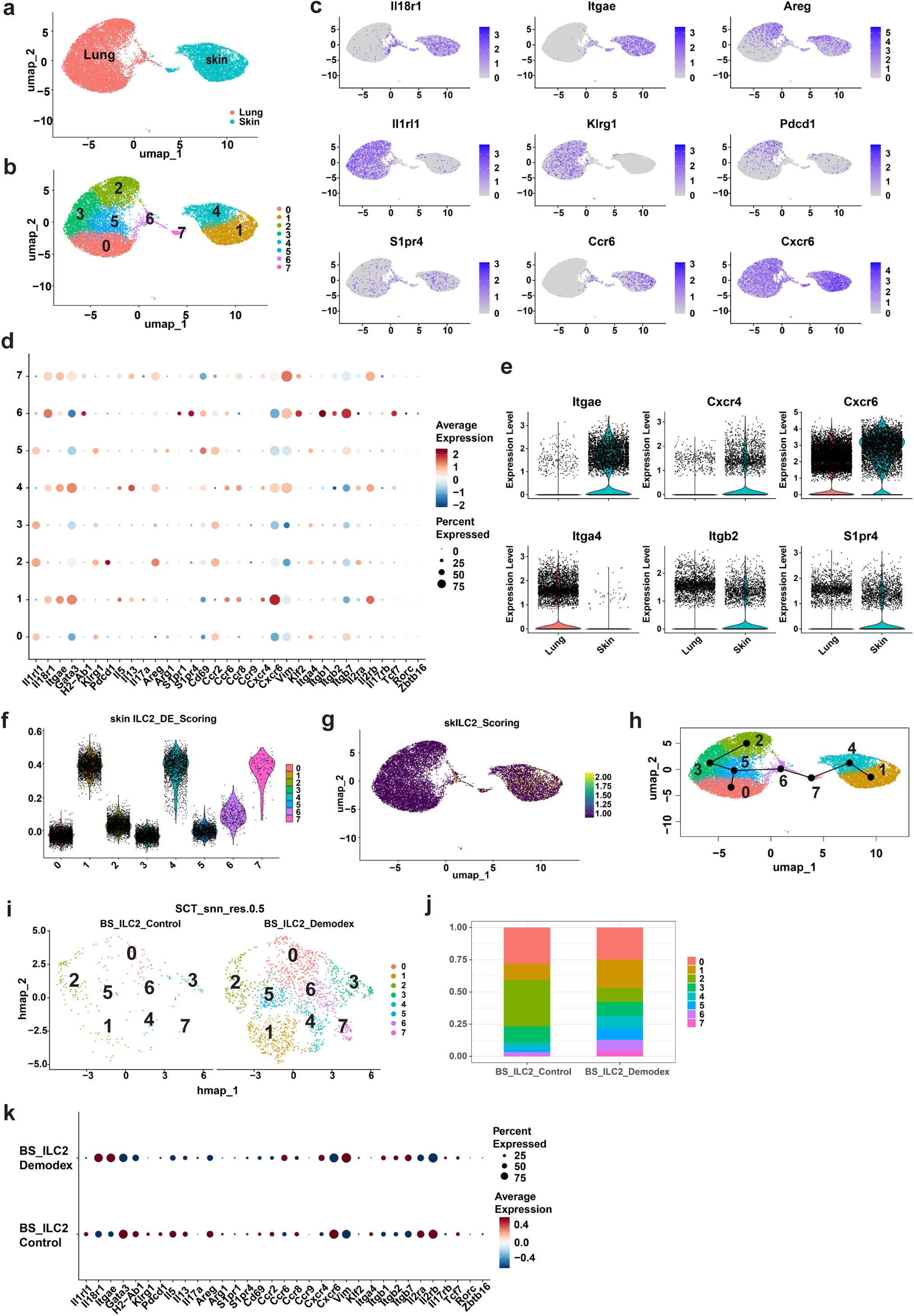
scRNA-seq analysis reveals migratory transcriptional signatures for skin-like ILCs. **a, b.** Sample tissue **(a)** and clusters distribution **(b)** for merged skin and lung Il5^RFP+^ ILC2s single cell RNA-seq (scRNA-seq) data set from GSE117568 (*Ricardo-Gonzalez et al., 2018*)^32^. **c.** UMAP for indicated genes for skin and lung ILC2s in (a)**. d.** Dot plot for indicated genes highlighting expression of transcripts associated with tissue-specific ILCs, effector function, migration and mobility, and tissue retention for each cluster in (**b**). **e.** Violin plot for indicated genes to compare expression between skin and lung ILC2s. **f.** Violin plot for skin ILC2 differential gene scoring for each cluster in **(b),** gene list in **Table S3. g.** UMAP for reparative skin-like ILC2 gene signature scoring. The featured skILC2 gene list that defined the score includes *Itgae, Il18r1, Ccr6, Ccr8, Il13, Il17a, Cxcr4, S1pr4, H2-Ab1, Klf2, Tcf7,* and *Itgb2.* **h.** Slingshot analysis for prediction of lineage differentiation. **i, j.** Sample **(i)** and clusters **(j)** distribution for ILC2s scRNA-seq from GSE197983 (*Ricardo-Gonzalez et al., 2022*)^28^ for control and Demodex mite-infected WT back skin. **k.** Dot plot for indicated genes transcription to compare ILC2s from control and Demodex mite-infected skin in **(i).**

### Parabiosis, adoptive transfer and skin tagging confirm the mobilization and lung retention of inflammatory skILC2s

To confirm that Demodex-infection promoted egress of skin ILC2s into the circulation and their subsequent recruitment to the lung, we performed parabiosis^36, 47^ between uninfected CD45.1/2 Il5^RFP^ (R5) and mite-infested CD45.2 IL4Rα^-/-^ mice. Consistent with our previous findings,^28^ skin ILC2s significantly expanded in the IL4Rα^-/-^ donor back skin and CD103^+^ skin-like ILC2s accumulated in the lungs (**Figure 5a**). The host mouse R5 lung also accumulated CD103^+^ skILC2s, the majority of which were host-derived, most likely triggered by new infection while joined with the mite-infested IL4Rα^-/-^donor; a fraction of these cells (10 - 15%), however, were recruited from the infested donor (**Figure 5b**). In contrast, in the mouse R5 liver, the percentage of skILC2s were equally distributed between donor (IL4Rα^-/-^) and host (R5), and not different than the distribution of circulating NK cells (**Figure 5b**). In both parabionts, lung ST2^+^ ILC2s and liver ILC1s remained host-derived reflecting their tissue residency. In contrast, donor-derived R5^+^ skILC2s contributed little to increases in host IL4Rα^-/-^ lungs and liver, which might be due to imbalanced expansion among the skin ILC2s between the WT and IL4Rα^-/-^ parabionts (**Figure 5a**), decreased cycling (Ki67^+^) of the re-circulating skILC2s in blood in IL4Rα^-/-^ mice and/or the prior occupation of available tissue niches in these mice (**Figure 1f, S1f**). Further, the majority of donor-derived skILC2s found in the IL4Rα^-/-^ lung remained intravascular (ivCD45^+^) while both donor and host-derived skILC2s in R5 lungs included ivCD45^+^ cells (**Figure 5c**), and consistent with the designation of CD103^+^ ST2^−^ skin-like ILC2s in non-skin tissues as ex-skILC2s.

**Figure 5.**
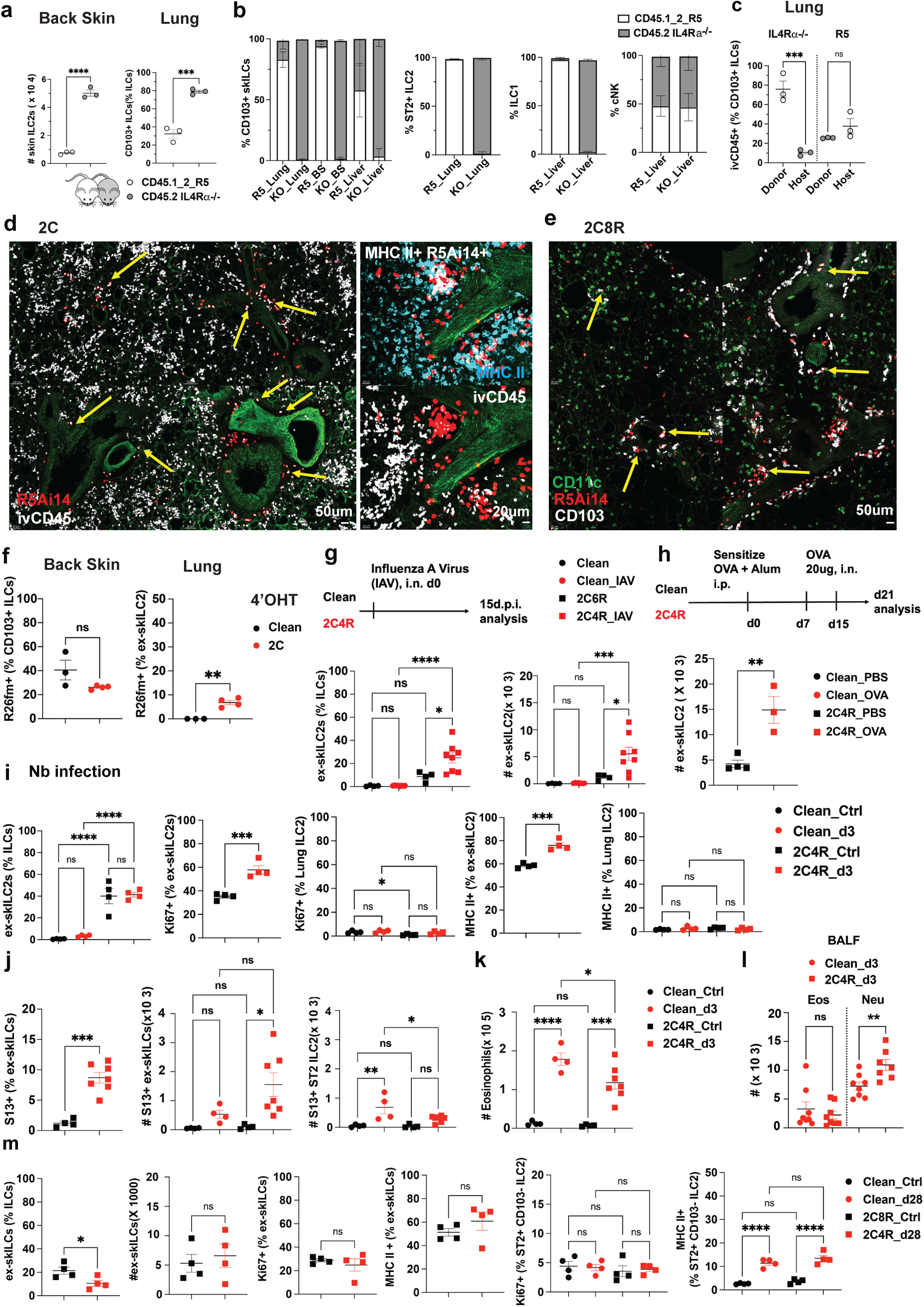
Recruited skin-like ILC2 establish residency in the lung and can respond to heterologous challenges. **a.** Parabiosis model of surgically linked 6-7wks old mice CD45.1/2 Il5^RFP^(R5) and CD45.2 IL4Rα^-/-^ (KO, Demodex-infested mice). Parabiosis pairs had a shared circulation for 4 weeks before tissue harvest. Comparison of CD103^+^ skin ILC2 (skILC2s) in back skin (left) and lungs (right). **b.** Percentage of CD103^+^ skILC2s, ST2^+^ ILC2, ILC1 and cNK in the indicated tissues of parabiotic mice. **c.** Donor vs Host-derived ivCD45^+^ CD103^+^ ILCs in the individual parabiont mouse. **d, e.** Immunofluorescence images from lung sections of R5Ai14 mice after 2 weeks (2C) or 2 weeks followed by 8 weeks of rest (2C8R) after Demodex exposure. Note the recently recruited ivCD45^+^ Il5^fm+^ (MHC II^+^) ILCs in 2C **(d)** and persistent CD103^+^ Il5^fm+^ ILCs in 2C8R **(e)**. **f.** Percentage of topical 4’OHT pre-labelled R26^tdTom+^ CD103^+^ skin ILC2s in clean and 2C back skin and lungs, respectively. **g.** Clean and 2C4R mice were infected with influenza A virus (IAV), and lungs were analyzed on d15 post-infection. Percentage (left) and cell count (right) of ex-skILC2 from the indicated groups. **h.** Clean and 2C4R mice were sensitized with OVA plus Alum at d0 intraperitoneal (i.p.) and then re-challenged with OVA intranasally (i.n.) at d7 and d15. Lungs were analyzed on d21, 6 days after the last intranasal administration of OVA. Graph of cell count of ex-skILC2 in indicated groups. **i.** Clean and 2C4R mice were infected with migratory helminth, *Nippostrongylus brasiliensis* (*Nb)*, and analyzed 3 days post-infection (3 d.p.i.). Graph show percentage of Ki67^+^ or MHC II^+^ ex-skILC2s or ST2^+^ ILC2s in lungs for indicated groups. **j**. Percentage (left) and cell count (middle) of IL-13 producing (Smart13^+^ (S13)) ex-skILC2s or ST2^+^ ILC2s (right) in lungs at d3 after *Nb* infection. **k.** Eosinophil counts in lungs from indicated groups as in (**j**). **l.** Cell counts for eosinophils and neutrophils in bronchoalveolar lavage fluid (BALF) for groups as in **(i). m.** Percentage and cell count of ex-skILC2s in lungs of indicated groups, and percentage of Ki67^+^ and MHC II^+^ for ex-skILC2s and CD103^-^ ST2^+^ ILC2 in lungs. *p< 0.05, **p< 0.01, ***p<0.001, ****p<0.0001 by unpaired t-test (**a, c, f, h**) or one-way ANOVA (**g, i-m**). n=3-8 biological replicates/group, representative of 1-2 independent experiments. Graphs depict mean ± SEM. Cell count and flow cytometry analysis are for the right lobe of the lung in f, g, h.

We performed a second parabiosis between neonatally infested, weaned and cleared adult CD45.1/2 IL4Rα^+/–^ (**Figure 2a**) and non-infected CD45.2 WT mice to assess the migratory capacity of lung-adapted ex-skILC2s. Most ex-skILC2s in the IL4Rα^+/–^ parabiont lung were ivCD45^−^ and consistent with lung residence (**Figure S6a**). In these parabionts conjoined in the absence of ongoing mite-induced skin inflammation, few ex-skILC2s were present in WT lung from either parabiont, or some remained ivCD45^+^, consistent with vascular residence despite the relatively high proportion of ivCD45^−^CD103^+^ ex-skILC2s in the lung of the previously infested pathobiont **(Figure S6b)**. As a second approach, we transferred total enriched CD45^+^ cells from the back skin of infected 2C R5Ai14 mice into IL7R^-/-^ mice, which significantly reduced competition by endogenous lymphocytes.^47^ Nearly all skin ILC2s from back skin of 2C mice expressed CD103 but not ST2, as expected (**Figure S6c**). Within two weeks of transfer, a fraction (∼15%) of the transferred skin-like CD103^+^ ILC2s gained expression of ST2 in IL7R^-/-^ lungs (**Figure S6d**), and the majority of these expressed CD69 (**Figure S6e**), consistent with tissue adaptation and residency. Together, these approaches suggest that 2 weeks of localized mite infection can provoke the appearance of blood and lymph node (**Figure S6f-h**) inflammatory (MHCII^hi^) ex-skILC2s (IL18R1^+^IL5^fm+^CD103^+^ST2^−^CD69^−^) that can give rise to IL18R1^+^IL5^fm+^CD103^+^ST2^+^ lung ILC2s, the majority of which gain CD69 and lose MHC II expression **(not shown)**, and consistent with adaptation, quiescence and tissue residence as predicted from lineage trajectories assessed using flowSOM (**Figure 3i, j, S3h, i, S4, S5**) and Slingshot (**Figure 4h**).^44, 45, 53^

As an additional approach, we used skin-localized tamoxifen (4’OHT)-induced genetic tagging^54^ to confirm that skin CD103^+^ ILC2s can migrate to the lung. We pre-labelled shaved back skin with 4’OHT daily for 5 days, and the labelling baseline was analyzed at d7, or followed by co-housing with Demodex mite-infested IL4Rα^-/-^ mice for 2 wks (**Figure S6i**) when 2C mice were analyzed (**Figure S6j,k**). Consistently, we found pre-labelled (R26^fm+^) CD103^+^ skin ILC2s recruited into lungs and skin-draining inguinal LNs (skdLN) but limited tissue adaptation by gain of ST2 (R26^fm+^ ST2^+^ CD103^+^ trILC2) in the lungs; lung resident ST2^+^ ILC2 were not labelled (**Figure 5f, S6l, m**). Although Demodex hair follicle infestation provoked skin ILC2 egress, this pathway is not provoked by general skin disruption since depilation resulted in skin ILC2 activation but not in the appearance of ex-skILC2s in the lung (**Figure S7a**).

### Re-activation of lung ex-skILC2s in the lungs upon immune perturbation

Immunohistochemistry of lung tissue confirmed the localization of newly recruited ivCD45^+^IL5^fm+^ and MHC II^+^IL5^fm+^ ex-skILC2s in lungs of 2C mice (**Figure 5d**) and persisting CD103^+^IL5^fm+^ ex-skILC2s in lungs of 2C8R mice (**Figure 5e**) within areas designated adventitial cuffs,^30^ thus sharing the microanatomical niche occupied by natural lung resident ILC2s. To assess the immune responsiveness of lung ex-skILC2s to a heterologous threat, we challenged WT or 2C4R mice with influenza A virus (IAV)^55^ and analyzed the lungs 15 days post-infection. While few ex-skILC2s are present in lungs from WT mice, the percentages and numbers of ex-skILC2s were increased among IAV-infected 2C4R mice (**Figure 5g**), confirming the expansion of ex-skILC2s in the new niche upon perturbation. We also sensitized mice using OVA-alum i.p. and analyzed the lungs 7 d after a second intranasal OVA challenge^56^ at d15 (**Figure 5h**). Again, both the percentages and numbers of ex-skILC2s were increased in mite-sensitized mice, confirming the capacity of migratory ex-skILC2s to respond to diverse lung challenges.

We next used *N. brasiliensis*^25, 33, 52^ to assess contributions of ex-skILC2s to this complex migratory infection that potently induces ILC2 activation, systemic migration of ILCPs and ILC2s, and their lung infiltration and differentiation. As early as day 3, which is prior to entry of gut migratory ILC2s^25, 52^ or de novo differentiation of ILCPs and immature ILCs,^33^ Ki67^+^MHCII^+^ ILC2s were increased in the lung in association with increased IL-13 expression (Smart13) in lungs of 2C4R mice as compared to non-mite-sensitized infected controls (**Figure 5i, j**). Curiously, eosinophils, although increased, were if anything lower and neutrophils were increased as compared to control mice, suggesting changes in the tissue immune response and consistent with the altered cytokine capacities (**Figure 5k, l**). To assess lasting effects in the lung associated with prior mite infestation, we compared *N. brasiliensis*-infected mice from both groups at day 28 after inflammation resolved and ex-skILC2s had assumed proliferation and activation profiles that differed little between the previously uninfected (Clean-Nb) and mite-infected animals (2C4R-Nb) (**Figure 5m**). Merging lung ILC2s from uninfected and *N. brasiliensis*-infected with previously mite-exposed uninfected and *N. brasiliensis*-infected mice (Clean, Clean-Nb, 2C8R, 2C4R-Nb, respectively) for UMAP- and flowSOM-mediated clustering based on multiparameter flow cytometric analysis revealed shared features, including differentiation of circulating and resident immature IL18R1^+^ ILCs as well as activation of ST2^+^ lung resident ILC2s and adaptation by KLRG-1^hi^ gut ILC2s, in agreement with prior studies,^33, 43, 52^ but additionally the appearance of activated ex-skILC2s, thus increasing immunologic diversity (**Figure 6a-c**). Indeed, flowSOM analysis supported the conclusion that *N. brasiliensis* infection drove ILCPs, inflammatory gut-derived ILC2s and resident lung ILC2s towards an inflammatory KLRG-1^+^ phenotype while activating ex-skILC2s that otherwise sustained characteristics of their tissue of origin (**Figure 6d, S7h**). The resulting lung profile revealed reductions in the overall population of activated KLRG-1^+^Ki67^+^MHC II^+^ ILC2s in *N. brasiliensis*-infected mice that had been previously exposed to Demodex (**Figure 6e**). Indeed, the percentage of ST2^+^ Th2 cells also decreased in these mice while CD103^+^ RORψt^+^ ψ8 T cells increased, and consistent with the type 2/3 mixed cytokine phenotype of ex-skILC2s and the alterations of neutrophils and eosinophils in the lungs (**Figure 6e, S7d, e)** and IgE in serum **(Figure S7b, c).**

**Figure 6.**
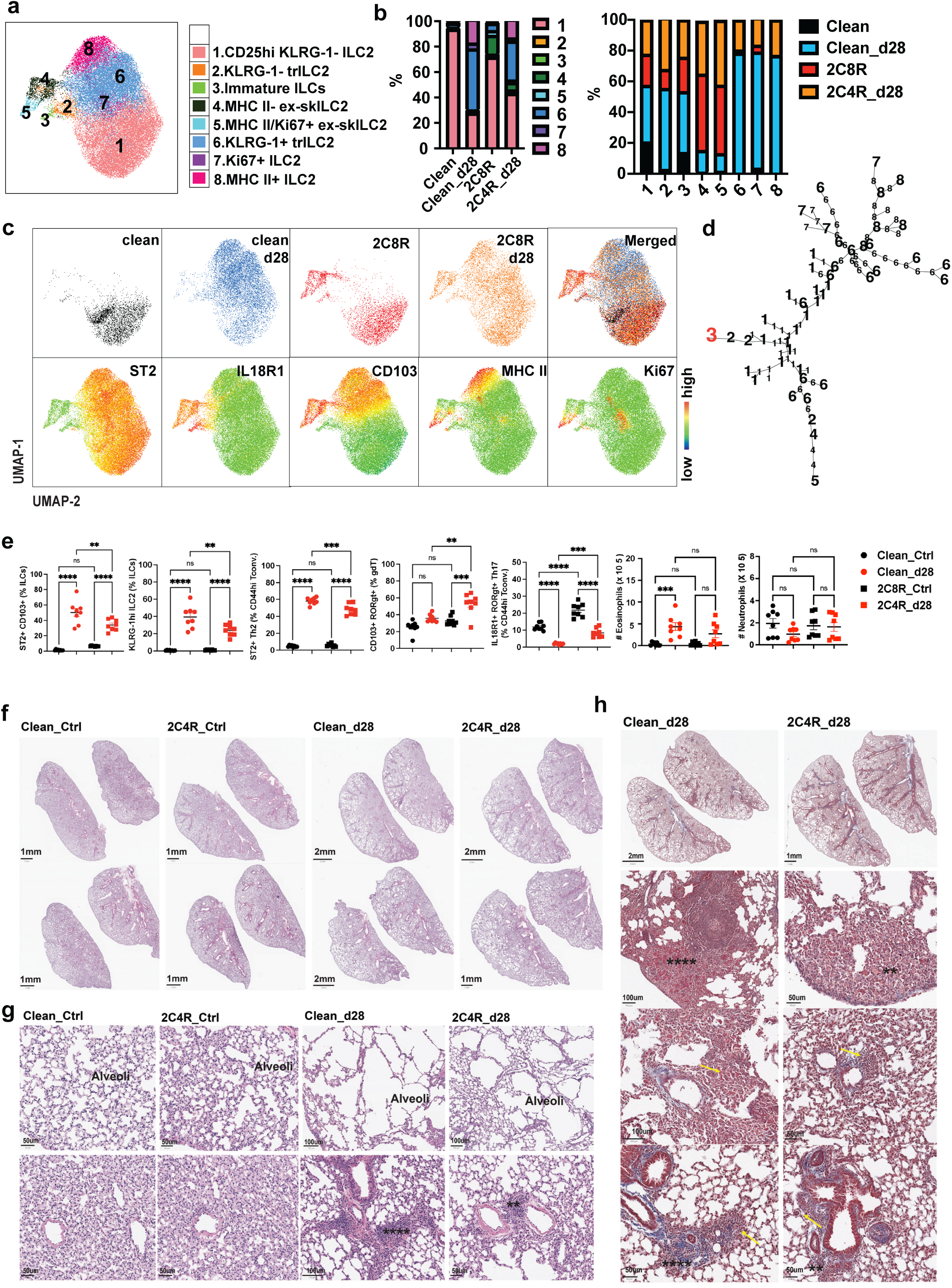
Demodex-experienced mice have a distinct ILC landscape in the lung and have an improved response to *Nb* infection. **a.** UMAP of merged ILCs populations analyzed by spectral flow cytometry for Clean and Demodex-experienced (2C4R) that were control or 28 days after infection with *Nb*. Clusters highlight the populations (pop) generated by flowSOM analysis. **b.** Clusters distribution and sample distribution for indicated groups as in (**a**). **c.** UMAP for the indicated experimental groups (top), and expression of the indicated markers (bottom). **d.** Predicted lineage tree by flowSOM analysis for *de novo* differentiation of immature IL18R1^+^ ST2^-^ CD103^-^ ILCs/ILC2P. Representation shows the predicted tissue adaptation of CD103^+^ ex-skILC2, and inflammatory differentiation of lung resident ILC2 (nILC2) and infiltrated inflammatory gut ILC2 (iILC2) at d28 after *Nb* infection. **e.** Percentage of ST2^+^ CD103^+^ trILC2, KLRG-1^hi^ iILC2, γδT, ST2^+^ Th2, RORγt^+^ Th17, and cell count for neutrophils and eosinophils in lungs from the indicated group at d28 after *Nb* infection. **f, g.** H&E section of whole fixed left lungs (**f**) and higher magnification (**g**) at d28 after *Nb* infection. Note the enlarged alveoli (empty space) and infiltrated leukocytes. Scale bar sizes as noted in the figures. Lung pathology scoring system: Pulmonary vascular and alveolar features, and bronchiole pathology was graded: normal, *mild, ** moderate, and ****severe (detail in **Table S1**). **g.** Masson’s Trichrome stain shows collagen deposition (blue) and fibrin (pink). *p< 0.05, **p< 0.01, ***p<0.001, ****p<0.0001 by one-way ANOVA (**e**). n=3-4 biological replicates/group, representative of 2 independent experiments. Graphs depict mean ± SEM.

To assess the impact that diminished overall inflammatory activation and enhanced early neutrophil influx might have on lung function, we compared lung pathology in the various uninfected and infected cohorts (**Table S1**). As assessed in 2C4R mice, mite-experienced *N. brasiliensis*-infected animals (2C4R-Nb) had less airway enlargement and diminished inflammatory cell infiltration into lung tissues (**Figure 6f-h**), with a trend towards reduced tissue transcripts for *Col1a1* and significantly less *Cthrc1* (encoding collagen triple helix repeat containing 1) and *Fn1* (fibronectin 1)^57^ (**Figure S7f**). Areg,^58, 59^ which has been associated with lung repair, was elevated at baseline in 2C4R mice (**Figure S7f**), and consistent with the reduced numbers of inflammatory ILC2s and Th2 cells, lower eosinophil infiltration and decreased total IgE (**Figure 6f, S7c, e**). Taken together, ex-skILC2s resident in lungs contributed to an altered inflammatory milieu that changed the immune and anatomic landscape of the subsequently perturbed lung (**Figure S7g**).

## Discussion

We describe activation of inflammatory skin ILC2s in response to the transient colonization of hair follicles of mice by ubiquitous Demodex mites. Although mites were cleared, migratory CD103^+^IL18R1^+^MHCII^+^ST2^-^ ex-skILC2s appeared in blood and within vascular beds in other tissues. In lung, but not other tissues examined, ex-skILC2s transitioned into parenchymal lung resident cells with a mixed lung (e.g., ST2^lo^) and skILC2 phenotype (e.g., CD103^+^) and mixed type 2-type 3 cytokine expression pattern that persisted for weeks after skin inflammation resolved. Ex-skILC2s responded earlier than resident lung ILC2s to diverse immune challenges and altered the cytokine and cellular landscape. Although several studies have called attention to immune gut-lung,^18, 25, 52, 60^ gut-skin and gut-brain circuits,^61, 62^ the physical migration of ILC2s educated post-birth from remote barrier tissues to the lung represents a cellular axis that imparts resilience to internal organs in concert with previously described metabolic and neural circuits.

Several aspects of the migration of ex-skILC2s were of note. First, activation and expansion in the skin was followed by blood entry and intravascular capture in organs, including the lung and liver, suggesting tethering by endothelial receptors while remaining labeled by anti-CD45 antibodies delivered via the bloodstream. Whether this reflects capture by distinct populations of endothelial cells within organs will require further study.^63, 64, 65, 66^ In contrast to other analyzed organs, lung-tethered ex-skILC2s persisted and slowly transitioned to tissue residence characterized by loss of iv CD45 antibody labeling and expression of markers shared with lung-resident ILC2s, such as ST2 and Arginase-l while several skin-associated markers, such as IL18R1 and CD103, continued to be expressed. The mechanisms for sustained tethering that allows adaptation to lung residence remain unknown, although we note prior observations supporting prolonged intravascular residence in lung for populations of activated immune cells^67, 68^ and the capacity for metastatic tumor cells to displace pericytes within lung endothelia to sustain access to intravascular nutrients while transitioning through metabolic adaptations required for parenchymal survival.^69^

Of interest, tethered, ex-skILC2s eventually accessed adventitial spaces where interstitial fluid collects within a network of stromal cells near broncho-vascular junctions identified as niches that support immune cells involved in surveillance, repair and defense of the lung.^30, 31^ How ex-skILC2s outcompete and persist in lung adventitial niches is unclear but the proportionate residence by these migratory cells was more successful when postnatal mice were infected as compared to adult mice. In the latter, greater proportions of surviving resident ILC2s matured from local lung and/or bone marrow precursors.^33^ The increased capacity for neonatal migratory ILC2s to adapt to the lung may reflect less competition for space during periods of early growth or better adaptation of perinatal ILC2s to developing tissues, or both,^33, 47, 70, 71^ and we speculate that better adaptation associated with early infestation may reflect temporal developmental windows that can contribute to the increasing prevalence of allergic lung disease in inverse relationship to public health standards as espoused by the hygiene hypothesis.^72^

Our studies additionally show that lung resident ex-skILC2s demonstrated earlier activation, earlier neutrophil accumulation in association with the increased capacity to generate IL-17, and potentially protection from lung damage, as previously demonstrated for migratory gut ILC2s.^25, 26, 52^ Together with the latter, these observations suggest the hypothesis that perinatal ILC2s, which undergo massive postbirth expansion in mice^47, 70^ can become educated at newly colonized barriers in the skin and gut by highly adapted eukaryotic colonizers like protists, helminths and hair follicle mites, and migrate to the lung, adapt to secondary residence, and contribute to resilience of the organ by accelerating the response to subsequent perturbation. The repositioning of educated ILC2s to internal sites may reflect the trajectory under natural conditions where exposure to diverse organisms in early perinatal life is more common. Loss of the latter may impact subsequent temporal plasticity as younger populations of precursor cells become replaced and postbirth tissue development slows. While further studies are required to assess the extension of our findings to humans, where ILC development occurs much earlier in utero, the identification of endogenous pathways that optimally drive migration and epigenetic changes among barrier ILC2s that facilitate internal tissue adaptation could suggest interventions that can overcome modern loss of commensal pressures that may have contributed to tissue resilience through evolution.

### Limitations of the study

Although we define migratory ex-skILC2s based on phenotype, transcriptomic, mRNA trajectory analysis, kinetics, and tamoxifen skin-limited fate-mapping, more definitive lineage tracking will require sophisticated genetic methods still being optimized. We used SPF mice challenged with natural commensals, including skin mites and worms, that likely colonize mice early in life in natural settings. As such, the migratory trajectory we document may occur normally, although it is possible that modern living standards have impacted these innate immune pathways, as discussed in the text. The mechanisms driving ILC2s from skin after Demodex infestation have not been ascertained. Finally, molecular cues that control lung vascular tethering and entry and adaptation to the lung adventitial niche as opposed to other tissues will require further study.

## Material/Methods

### Mice

Wild-type (C57BL/6J; Stock 000664; B6 CD45.1; Stock 002014) mice were purchased from Jackson Laboratories. IL4Rα^-/-^, IL4Rα^+/-^ x Arg1^YFP^ x Il5^RFP^, Arg1^YFP^ x Il5^RFP^, Il5^cre^ x R26Ai14 on C57BL/6 backgrounds were bred and maintained as described.^28, 29, 40, 43^ Homozygous Rosa26^CreERT2^ (Stock 008463) and R26Ai14 (Stock 007914) were purchased from Jackson Laboratory and intercrossed to generate double heteryzogous mice for these alleles. Sex-matched mice aged 7-14 weeks were used in all experiments unless otherwise indicated. Mice were maintained under specific pathogen-free conditions. All animal procedures were approved by the UCSF Institutional Animal Care and Use Committee.

### Demodex-mite co-housing for infestation

To infest mice with Demodex, we co-housed female mice with designated alleles at ages 4-6 wks by co-housing with a mite-infested IL4Ra^-/-^ female mouse.^28^ Infection occurred naturally during co-housing over 2 wks; newly infested recipient mice were designated 2C. Mice were separated to allow mite clearance (typically over ∼14-21 days) and analyzed after 4 or 8 wks, termed 2C4R and 2C8R, respectively.

### Intravenous anti-CD45 labeling

To label intravascular leukocytes, 2 μg of anti-mouse CD45 antibody diluted in 200 μl PBS were injected via retroorbital vein after isoflurane anesthesia.^37^ Mice were euthanized within 5 min after antibody injection.

### Parabiosis

Parabiosis was performed as described,^36, 47^ resulting in 40%–60% mixing of circulating blood cells by 2 wks. Briefly, pairs of mice were joined by suturing mirror-image peritoneal openings, elbow and knee joints, followed by skin stapling as necessary (9 mm autoclip, Clay Adams). Post-op pain was managed using buprenorphine. Parabiosis pairs were euthanized 3-4 wks after surgery and blood and tissues (lung, liver, back skin) were collected for processing and flow cytometric analyses.

### Depilation experiment

We performed depilation experiments as reported.^28^ Briefly, mice were anesthetized with isoflurane inhalation. Under anesthesia, the dorsal surface (back) hair of the animals was shaved down to the level of skin with an electric razor. A thin coat (1 fingertip unit) of Nair depilatory cream (Nivea) was applied to the shaved region for a period of 30 seconds before wiping clean. Mice were analyzed at d14 after depilation was performed.

### Administration of 4’OHT

In experiments using topical 4’OHT (Sigma-Aldrich), 5 mg of 4’OHT was dissolved in 10 ml of acetone (Sigma-Aldrich) to prepare a stock solution at 500 μg/ml. 4’OHT was further diluted with acetone to generate working concentrations of 20 μg/ml (1:25 dilution). 120 μl of 4’OHT/acetone at working concentration was then applied to anesthetized mice in a dropwise manner to the shaved dorsal back skin of mice to cover the entire shaved skin surface (no depilation is performed).^54^ Given the relatively high vapor pressure of acetone, there is no residual liquid to remove from the skin, and the mice are returned to their cage. This procedure is repeated for 5 consecutive days, and mice were analyzed 2 days after the last treatment or co-housed with a mite-infested IL4Rα^-/-^ female mouse and analyzed at the indicated time points.

### Helminth infection

Mice were injected subcutaneously (s.c.) with 500 L3 *Nippostrongylus brasiliensis (Nb)* larvae prepared as described.^73^ Briefly, *Nb* was maintained by passage in rats with stool used to purify larvae by mixing with water and activated charcoal before culture in Petri dishes at 25°C for up to 4 wks in a humidified chamber. L3 *Nb* were separated using a Baermann apparatus after 7 d culture. L3 larvae were extensively washed with PBS prior to injection and mice were analyzed at designated periods after infection.

### OVA sensitization

Matched uninfected or mite-exposed 2C4R mice were immunized with 20 μg ovalbumin (OVA, Sigma, A5503) plus 2 mg alum (Thermo Fisher, 77161) in 200 ul PBS by i.p injection.^73^ Mice were re-challenged intranasally with 20 μg OVA in 40 ul PBS at d 7 and d 14 and analyzed on d 21.

### Influenza A infection

Mice were challenged by intranasal aspiration of 100 plaque-forming units (pfu) of influenza A virus A/PR/8/34 as described.^74^

### Tissue dissociation and cell isolation

Lung,^32, 33^ skin^28^ and liver^75^ tissue single-cell suspensions were prepared as described. Briefly, lungs were dispersed using the automated tissue dissociator (GentleMACS; MiltenyiBiotec) following 37°C incubation on the shaker at 120 rpm for 20 min in 5 ml RPMI 1640 digestion buffer (10 mM HEPES, 2% FBS, 0.1 mg/ml Liberase TM, 0.1 mg/ml DNase I); for combined myeloid cells analysis, lungs were physically dissociated using scissors and incubated for 40 min in 3 ml RPMI 1640 digestion buffer. Digestion was stopped using cold 20 ml FACS Buffer (PBS supplemented with 3% FBS). After subcutaneous fat was removed, back skin single-cell suspensions were prepared as described^35^ Briefly, skin tissues were minced in RPMI-1640 with 5% FBS, transferred to C tubes (Miltenyi Biotec) containing 5 ml RPMI-1640 (Sigma R8758) supplemented with Liberase TM (0.25 mg/ml) and DNase I (0.1 mg/ml). Samples were shaken at 180 rpm for 1.5 h at 37 °C and dispersed using the automated tissue dissociator running program C. Cells were filtered through 70 μm strainer (Falcon, 3522350) and washed and collected in FACS Buffer. For liver, after tissue digestion, lymphocytes were enriched in 40% vs 80% Percoll gradients at 2500 rpm for 20 min at 20 °C. The interface was washed with FACS buffer. For purification of bone marrow cells, the bone was opened, the marrow was flushed with cold FACS buffer, pipetted to break up aggregates and centrifuged at 500 x G for 5 min. Peripheral blood was collected directly from the left ventricle. Red blood cells were lysed using BD Pharm Lyse lysing solution (BD, 555899) before further analysis.

### Flow cytometric analysis

Cell suspensions prepared as above were incubated with Fc block (2.4G2, Bioxcell) and Fixable Viability Dye, and stained with fluorophore-conjugated antibodies against CD3ε, CD5, CD19, Gr-1, F4/80, CD11b, CD11c, NK1.1, TCRβ, TCRγδ, FcεR1, TER-119, Thy1.2, CD127/IL7Ra, CD25, IL25R/IL17RB, CD218a/IL18Ra, ST2, KLRG-1, PD1/CD279, α4β7, C-kit, CD45.1, CD45.2, CD45, NK1.1, I-A/I-E, CD44, CD103, CD69, RORγt, Gata3, Foxp3, and Ki67. Lineage (Lin) positive cells were stained with a cocktail of biotin-labeled antibodies against CD3ε, CD5, Gr-1, CD19, TCRβ, CD11b, CD11c, F4/80, TER-119 and FcεR1, followed by staining with Streptavidin (V500, BD) and secondary antibodies to allow selection of cells negative for CD4, CD8a, TCRγδ and NK1.1. For intracellular cytokine staining, tissue preparations were cleared using a Percoll gradient (Sigma, P4937). Enriched lung cells were cultured in RPMI-1640 supplemented with 10% FBS, and re-stimulated with 50 ng/ml phorbol myristate acetate (PMA) and 500 ng/ml ionomycin in the presence of 1 μg/ml monensin with 1 μg/ml Brefeldin A for 3.5 h. Intracellular staining for CD3ε, IL5, IL13, IL17A, Ki67, Gata3 and RORγt was performed using Foxp3/Transcription Factor Staining Buffer Set (eBioscience, 00-5523-00). For flow cytometric analysis, cells were analyzed on a 5 laser Cytek Aurora cytometer and analyzed with FlowJo software using tSNE, UMAP and flowSOM analyses where designated.

### Single-cell RNAseq data re-analysis

Single-cell RNA-seq data were analyzed using Seurat version 5 with standard workflows. Il5^RFP+^ lung and skin ILC2s (**GSE117568**)^32^ were first merged by Seurat function (JoinLayers) for tissue-specific ILC2 transcriptional analysis, while skin ILC2 data from Clean and Demodex-infested mites (**GSE197983**)^28^ were from WT skin Thy1_pos data set function (RunHarmony) used for batch correction. After quality control, viable cells were narrowed by excluding cells with UMI counts lower than 200 and above 2500 or if containing more than 5% mitochondrial transcripts. The 2000 most variable genes used for the anchoring process were used for downstream analysis to calculate principal components after log-normalization and scaling. Principle component analysis (PCA) was used for dimensionality reduction and to visualize a uniform manifold approximation and projection (UMAP) for identified clusters. P-values comparing gene expression of clusters and samples were calculated using FindMarker function in Seurat. Gene list for **Table S3** was generated using an adjusted p-value cutoff <0.05 and expression in the indicated cluster cutoff >50%. Gene signature scores were calculated using the AddModuleScore function in Seurat. Gene lists to calculate ILC2 skin scores and core gene signatures were obtained from top100 differential gene expression (**Table S3)** and gene lists, respectively. Trajectories were predicted using Slingshot with standard settings.

### Tissue imaging

Right lung lobes and inguinal lymph nodes from B6, Il5^RFP^ and Il5^cre^ x R26Ai14 were fixed in 1% PFA at 4°C overnight. Tissues were washed with PBS, incubated in 30% sucrose at 4 °C for 8-24 h, embedded in Optimal Cutting Temperature Compound (Tissue-Tek) and stored at –80 °C. Tissue sections (50 μM) were cut using a Leica Cryostat (Leica). Sections were blocked with 0.1%Triton X-100, 2% BSA, and 2% normal mouse serum in PBS at RT for 1 hr. Primary antibodies included anti-Mouse MHC II Pacific Blue or anti-Mouse EpCAM BV421 and anti-mouse CD11c AF488 or/and CD3ε AF488 as indicated. Antibodies (BioLegend) were diluted (1:200) in staining buffer (PBS, 0.1% Triton X-100, 2% BSA, 2% normal mouse serum) and incubated at 4°C overnight. Goat anti-mouse CD103 (R&D, AF1990-SP) and Living Colors DsRed Polyclonal antibody (rabbit, 1:300, Takara Bio #632496) as designated were used at 1:400, followed by donkey anti-goat AF555 and donkey anti-rabbit AF647 at 1:800 at 4 °C for 8-12 h. Slides were mounted in Mounting Media (Invitrogen, 00-4958-02). Images were taken using a Nikon A1R-TIRF confocal scanning microscope with Plan Apo λ 20X/0.75NA air objective running NIS Elements software. Image analysis was performed with Imaris (Oxford Instruments).

### Tissue histology

Lung tissue was fixed in 4% neutral buffered formalin and submitted for paraffin embedding, hematoxylin & eosin and Masson’s Trichrome staining to HistoWiz (Long Island City, NY). Sections were analyzed using QPath open software.^76^

### RNA preparation and RT-PCR

Lung tissues were collected and stored in RNAlater (Fischer Scientific, AM7020) at –80 °C until processed. RNA extraction was performed using RNeasy Micro Plus kit (Qiagen) per the manufacturer’s protocol. For quantitative reverse transcriptase PCR analyses, RNA was reverse transcribed using the SuperScript VILO cDNA synthesis kit (ThermoFisher) per the manufacturer’s instructions. The cDNA was used to test expression of select genes using Power SYBR Green PCR master mix (ThermoFisher) in a StepOnePlus cycler (Applied Biosystems). Primer sets from PrimerBank for *Col1a1, Cthrc1, Spp1, Fn1, Arg1,* and *Areg* are listed in **Table S2**. Gene expression was normalized to *Rps17* using the ΔCt method.

## Statistical analysis

Experiments were performed using randomly assigned mice without investigator blinding. All data points and n values reflect biological (individual mice) replicates. Experiments were pooled whenever possible and data were analyzed using Prism 10 (GraphPad Software) by comparison of means using unpaired t*-*test or one-way ANOVA for multiple comparison as designated in figure legends. Data in figures represent mean ± SEM unless indicated otherwise.

## Author contributions

Conceptualization: M.L., R.R.R.-G., R.M.L.

Methodology and investigation: M.L., F.L.P, K.K., L.H.Q., T.R., L.T., H.E.L., R.R.R.-G.

Mouse Models: Parabiosis: L.H.Q.; Helminth infection: H.E.L.; Influenza A infection, Depilation, 4’OHT: R.R.R.-G.

Formal analysis, visualization and data curation: M.L. (FACS analysis, UMAP, flowSOM/Seurat/ Confocal Imaging/H&E analysis); F.L.P. (Seurat/Harmony); K.K. (Seurat Scoring/Slingshot); T.R & L.T. (real-time PCR)

Writing - original draft: M.L.

Writing - review and editing: M.L., R.R.R.-G., R.M.L., all authors

Funding acquisition and supervision: R.R.R.-G., R.M.L.

## Acknowledgements

We thank the UCSF Animal Facility for mice husbandry and technical assistance. We thank the support from UCSF Flow Core, and UCSF Imaging Core. We thank Dr. Maya Kotas for sharing Influenza A PR8 virus and Dr. Ari Molofsky for the R5-Ai14 mouse line. We thank Ouyang Ye (Gasteiger Lab) for technical assistance.

This work was supported by the National Institutes of Health (P01HL107202 to R.M.L and K08AR075880 to R.R.R.-G.), Howard Hughes Medical Institute (R.M.L.), Biohub San Francisco (R.R.R.-G.) and Mount Zion Health Fund (R.R.R.-G.).

## Supplemental information

**Figure S1.**
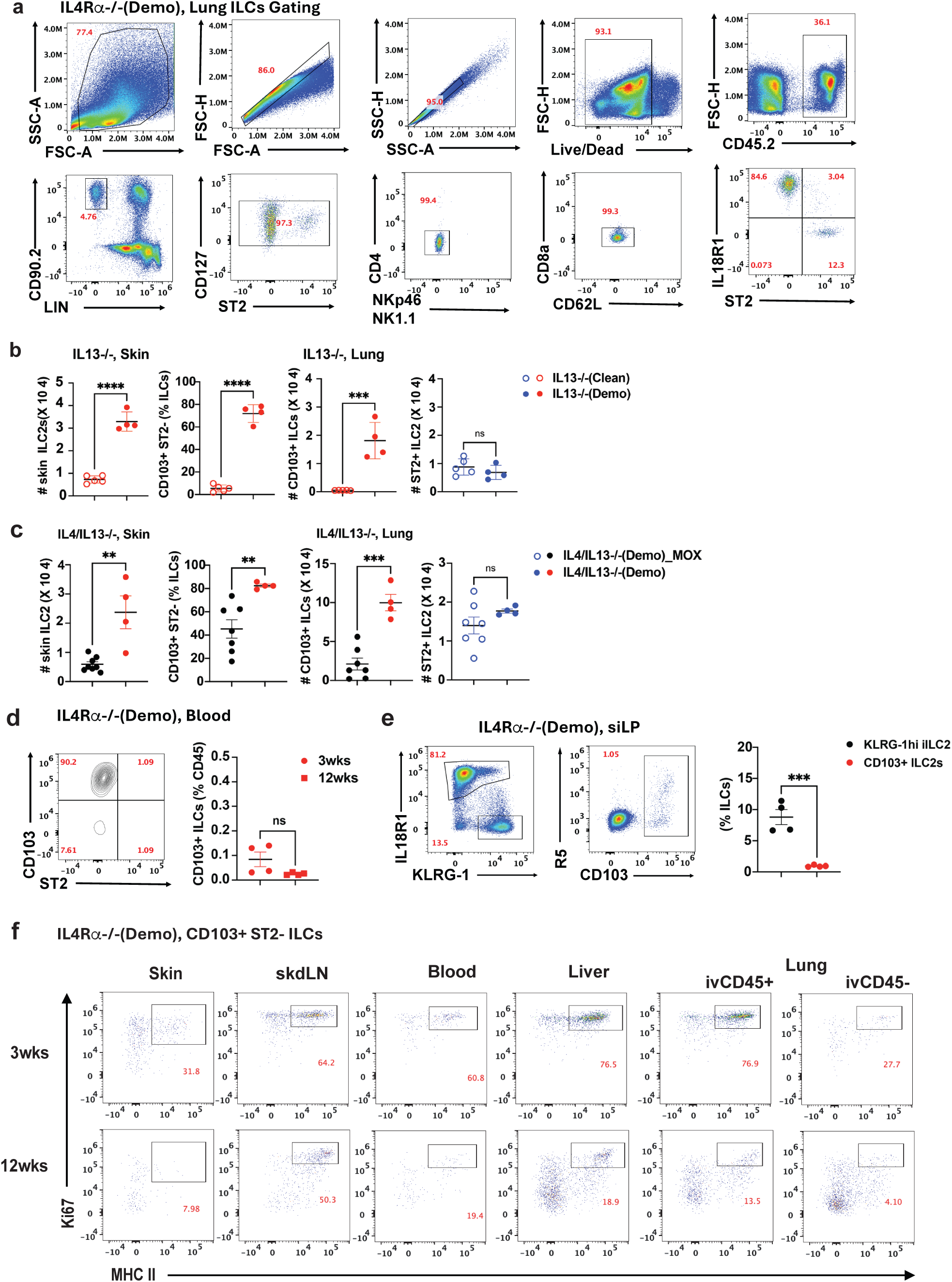
Skin-like ILC2s in Demodex mite-infected IL4Rα^-/-^ mice enter circulation and are present in multiple organs. **a.** Gating strategy for CD127^+^ ILCs in IL4Rα^-/-^ (Demo) lung. **b.** Cell count of CD103^+^ ILCs in skin, percentage and cell count of CD103^+^ skin-like ILCs in lungs, and cell count of ST2^+^ ILC2 in lungs from clean and Demodex mite infested IL13KO mice. **c.** Percentage and cell count for ILCs in skin and lungs from control (mite infested) or moxidectin/imidacloprid (MOX, cleared of Demodex mites) IL4/IL13DKO mice. **d.** FACS plot and percentage of skin-like ILC2 (Lin^-^CD90.2^+^IL7R^+^IL18R1^+^CD103^+^ST2^-^) in IL4Rα^-/-^ (Demo) peripheral blood. **e.** Gating for KLRG-1^hi^ ILC2 and CD103^+^ skin-like ILC2 in IL4Rα^-/-^ (Demo) small intestine lamina propria (siLP). **f.** FACS plot for Ki67 and MHC II expression on CD103^+^ ST2^-^ ILCs from indicated tissues of IL4Rα^-/-^ (Demo) mice at age of 3wks and 12wks. ***p<0.001 by unpaired t-test. n=3-4 biological replicates/group, representative of 2-3 independent experiments (**c, e**) or pooled from 3 experiments (**f**).

**Figure S2.**
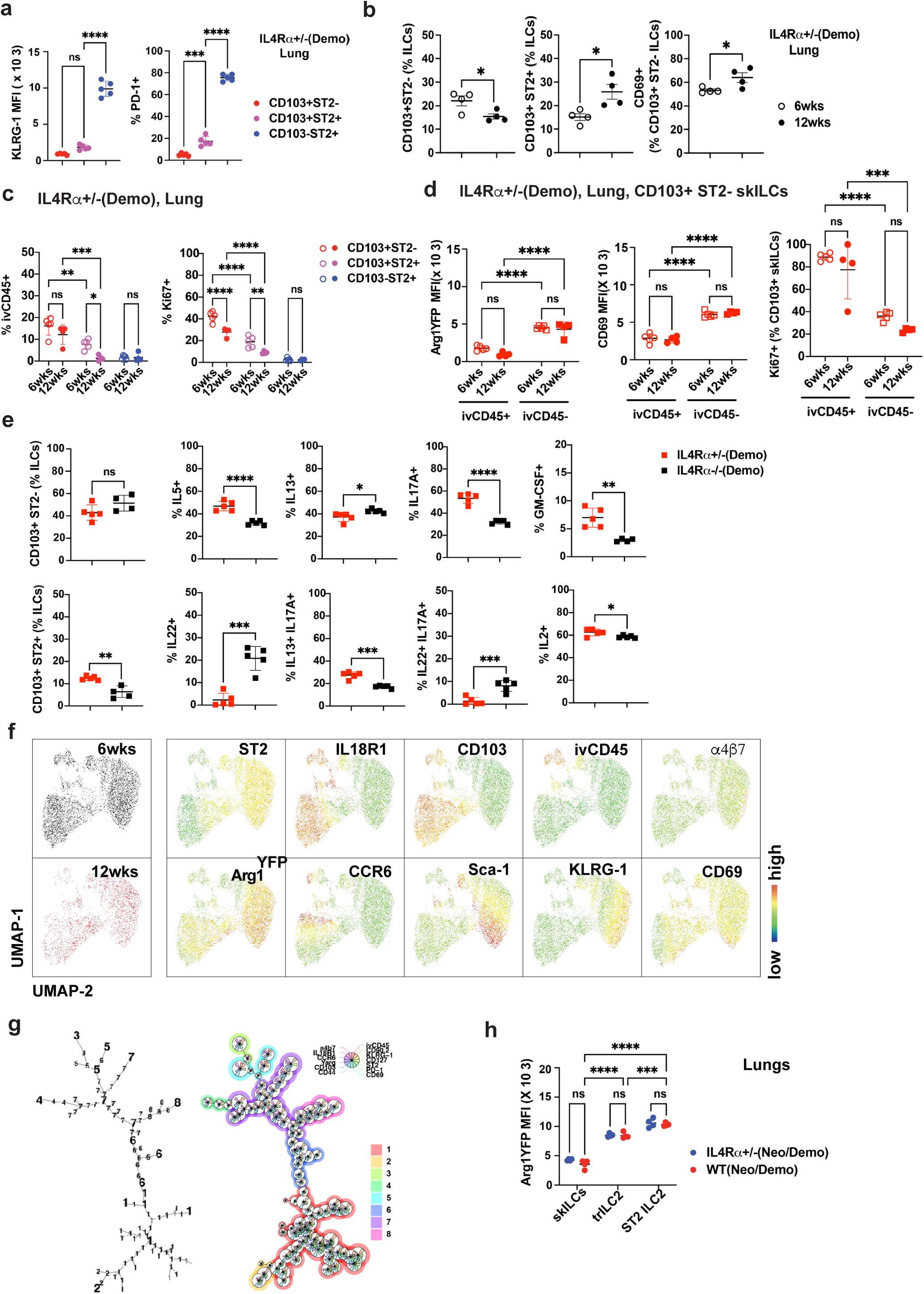
Heterogeneity of lung ILC2s from IL4Rα^+/–^ (Demo) mice. **a.** KLRG-1 MFI and percentage of PD-1^+^ for indicated ILC subsets, CD103^+^ ST2^-^, CD103^+^ ST2^+^ and CD103^-^ ST2^+^ ILC2s in IL4Rα^+/–^(Demo) lungs. **b.** Percentage of CD103^+^ ST2^-^ and CD103^+^ ST2^+^ ILC2s, and CD69^+^ CD103^+^ ST2^-^ skin-like ILCs in IL4Rα^+/–^ (Demo) lungs at age of 6wks and 12wks. **c.** Percentage of lung ILC2 subsets as in **(a)** that are ivCD45^+^ and Ki67^+^ at 6 and 12wks of age, respectively. **d.** Comparison of Arg1^YFP^ MFI, CD69 MFI, and percentage of Ki67^+^ cells of ivCD45^+^ versus ivCD45^-^ for indicated lung ILCs subsets. **e.** Percentage of CD103^+^ ST2^-^ and CD103^+^ ST2^+^ in Demodex-mite infested IL4Rα^-/-^ (black) and IL4Rα^+/–^ (red) lungs (left panels), cytokine profiling (right panels) for CD103^+^ ILCs upon *in vitro* PMA/Ionomycin re-stimulation. **f.** UMAP and indicated markers for merged ILCs from IL4Rα^+/–^ (Demo) lungs at 6 and 12wks of age. **g.** Arg1^YFP^ MFI for indicated lung ILC2 subsets for Neo/Demo WT (red) and IL4Rα^+/–^ (blue) mice. *p< 0.05, **p< 0.01, ***p<0.001, ****p<0.0001 by unpaired t-test. n=3-5 biological replicates/group, representative of 1-2 independent experiments.

**Figure S3.**
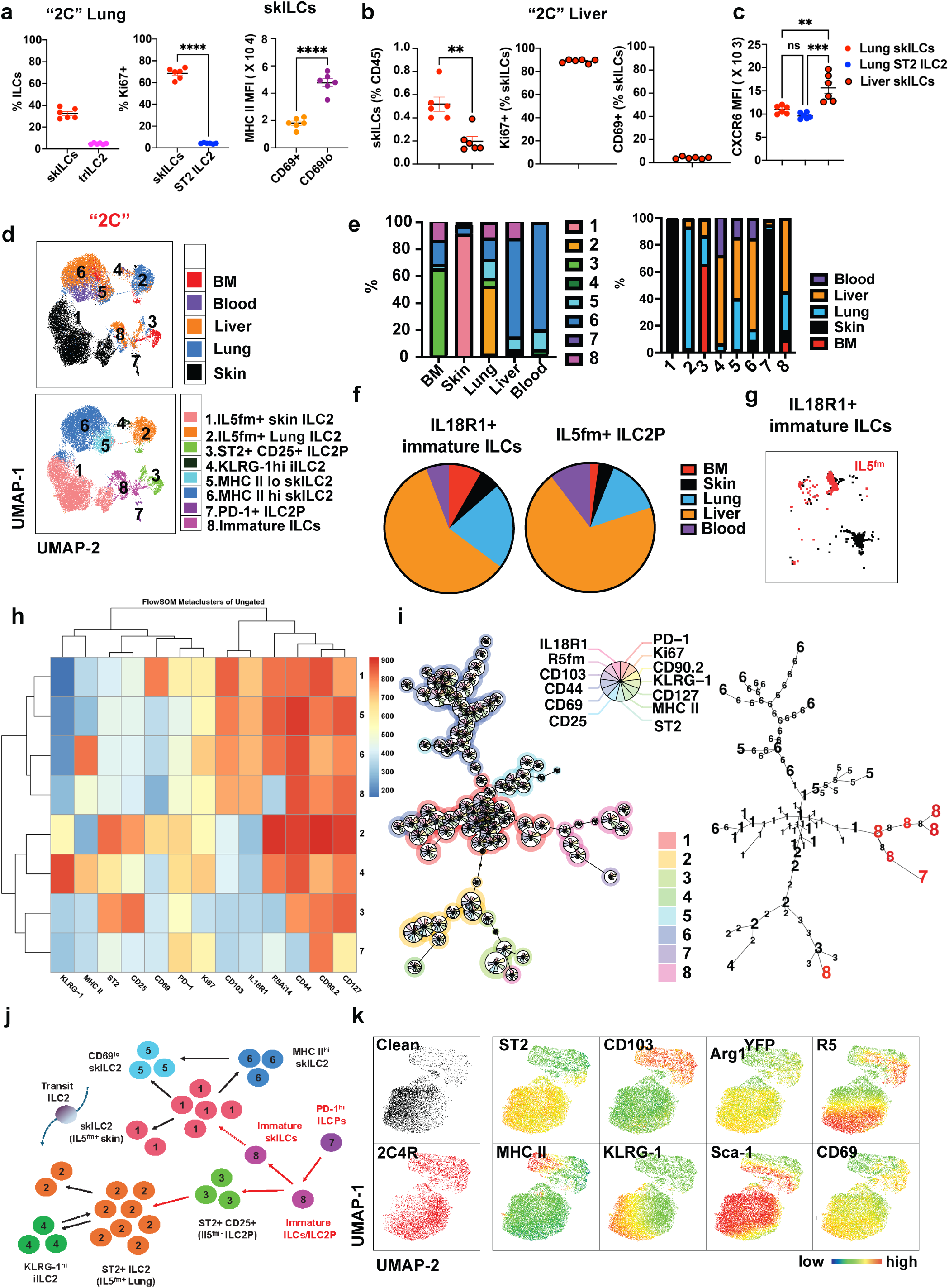
Adult mice exposed to Demodex mites establish a distinct lung ILC landscape. **a.** percentage of ST2^-^ CD103^+^ and ST2^+^ CD103^+^ ILC2s, percentage of Ki67^+^ for CD103^+^ skILCs and ST2^+^ ILC2, and MHCII MFI for CD69^+^ versus CD69^lo^ CD103^+^ skILCs in 2C lung. **b.** Comparison of skILCs in 2C lung and liver (left), percentage of Ki67^+^ and CD69^+^ for skILCs in 2C liver. **c.** CXCR6 MFI for lung skILCs, lung ST2^+^ nILC2, and liver skILCs in 2C mice**. d.** UMAP of spectral flow cytometry of the merged ILCs from 2C mice bone marrow, liver, lung, back skin and peripheral blood (top) and UMAP highlighting the individual cluster populations (pop) identified by flowSOM analysis of merged ILCs (bottom). **e.** Clusters distribution among samples (left) and tissue origin distribution within clusters (right) as in **(d). f, g.** Pie charts for IL18R1^+^ ST2^-^ CD103^-^ immature ILCs and IL18R1^+^ ST2^-^ CD103^-^ IL5^fm+^ ILC2Ps in peripheral tissues of 2C mice **(f)** and UMAP for immature ILCs and IL5^fm+^ ILC2Ps **(g). h.** Heatmap for distinct ILC clusters (pop 1 to 8) and their expression of the indicated markers by flowSOM analysis. **i.** Lineage trees for merged tissue-specific ILC2s from 2C mice, represented with their population number (left) and population markers (right). **j.** Summary for predicted lineage tree and *de novo* differentiation of immature ILCs and/or IL5^fm+^ ILC2Ps and tissue adaptation of skin-like ILC2s in lungs. **k.** UMAP for clean versus 2C4R lung ILCs and the expression of selected markers by flow cytometry. *p< 0.05, **p< 0.01, ***p<0.001, ****p<0.0001 by unpaired t-test. n=3-6 biological replicates/group, representative of 2-3 independent experiments. Merged tissue-specific ILCs for UMAP and flowSOM analysis are representative n=2 biological replicates.

**Figure S4.**
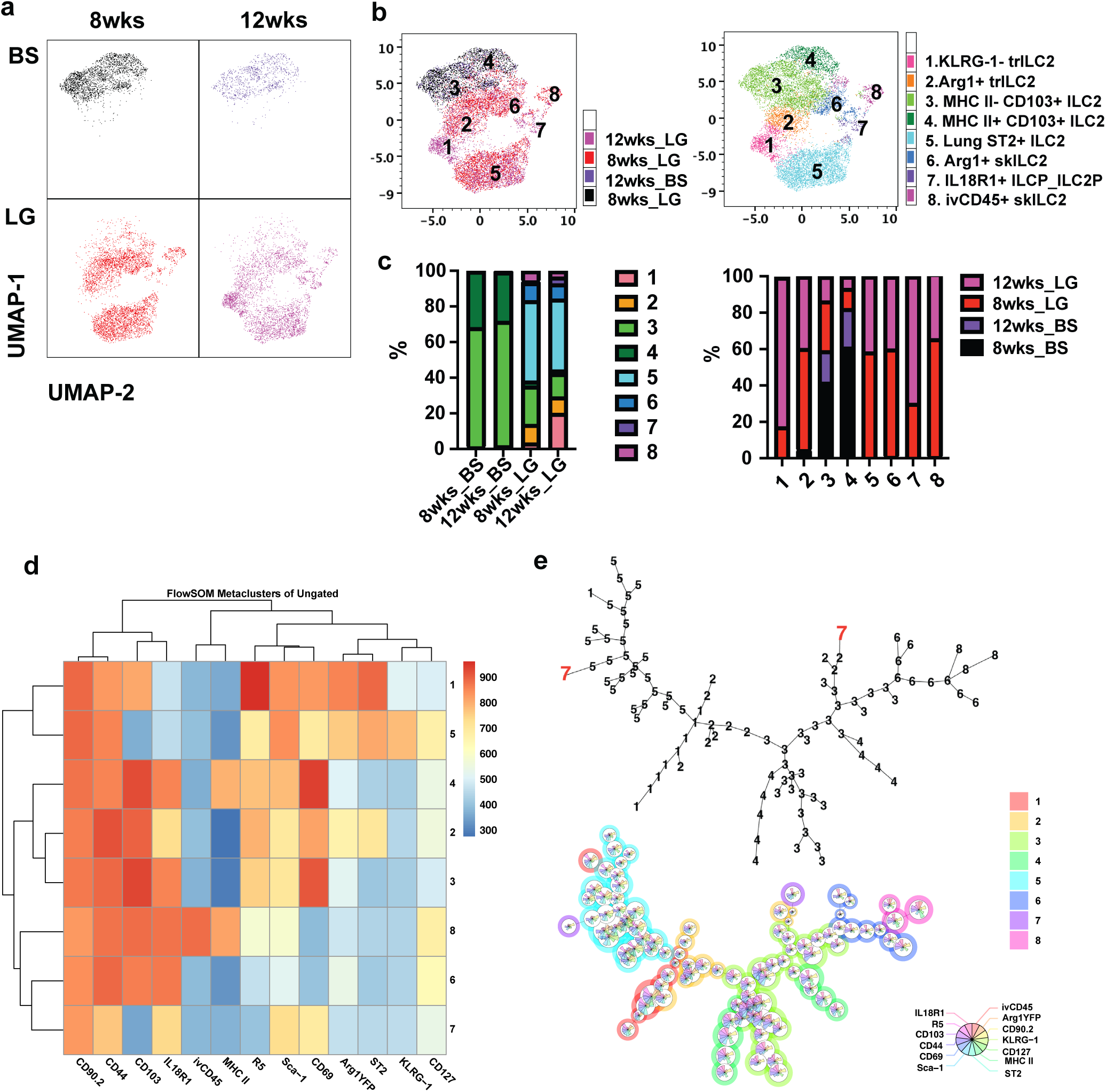
Demodex pre-exposure mobilizes inflammatory CD103^+^ skin ILC2 that adapt to the lung environment. **a, b.** UMAP showing individual samples distribution (**a**), merged UMAP (**b**) samples (left) and ILC clusters distribution (right) identified by flowSOM analysis of merged ILCs in Neo/Demo IL4Rα^+/–^ lungs (LG) and back skin (BS) at age of 8wks and 12wks. **c.** Clusters (pop) distribution among samples (left) and sample distribution within clusters (right) in (**a**)**. d.** Heatmap for selected expression of surface markers by the individual ILC clusters (population 1 to 8) by flowSOM analysis. **e.** Lineage trees for merged tissue-specific ILC2s from IL4Rα^+/–^(Neo/Demo) mice, represented by population number (top) and by their expression of markers (bottom). Merged tissue-specific ILCs for UMAP and flowSOM analysis, representative n=2 biological replicates.

**Figure S5.**
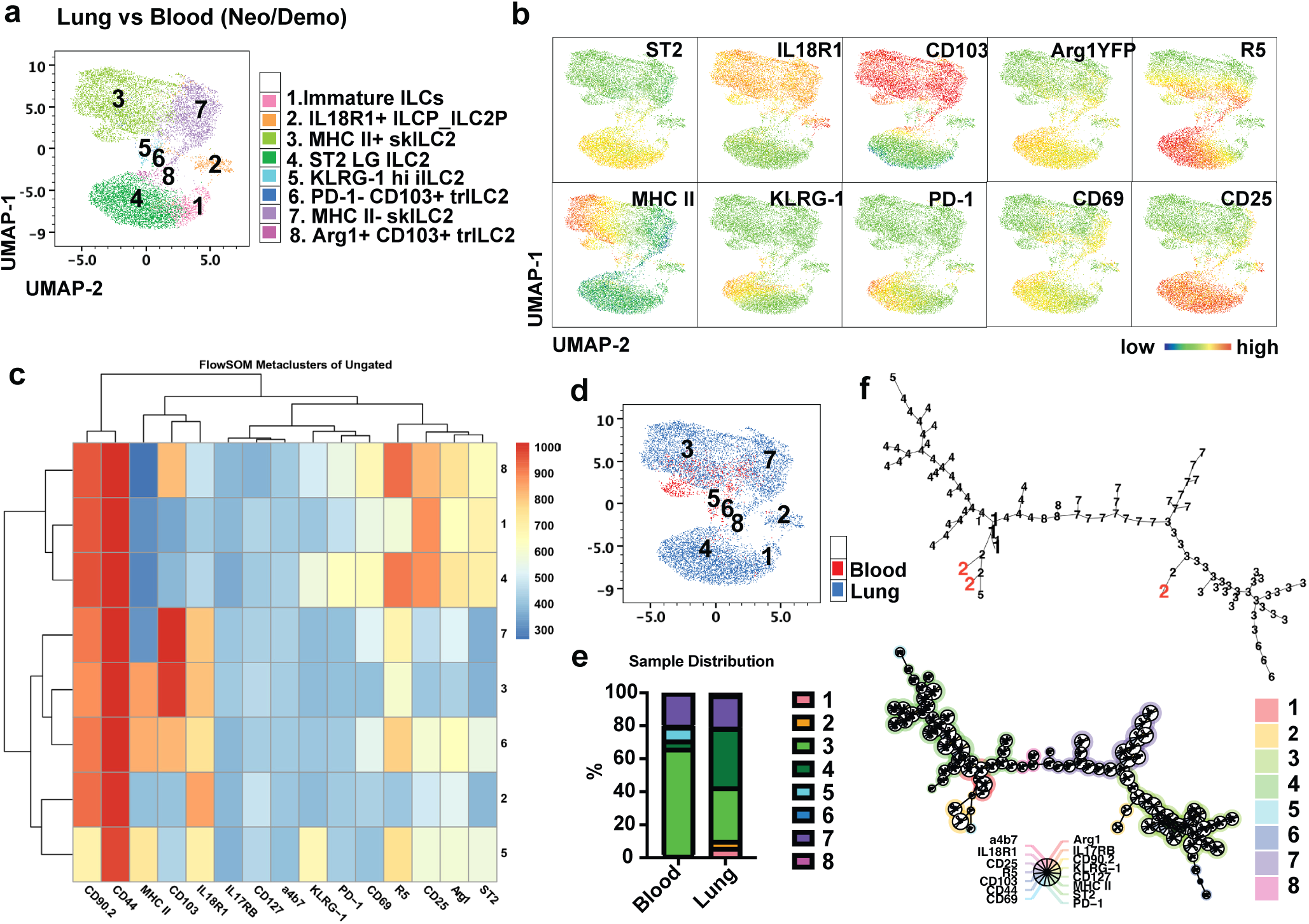
Demodex pre-exposure mobilizes circulating skin-like ILCs that adapt to the lung environment. **a.** UMAP generated from spectral flow cytometry data of merged lung ILCs and peripheral blood Il5^RFP+^ ILCs (R5) of mice exposed to Demodex during the neonatal period. **b.** UMAP of surface markers expressed by ILCs. **c.** Heatmap for the indicated markers by individual clusters of ILC populations. **d, e.** UMAP representation of samples **(d)** and distribution of populations within each sample **(e)**. **f.** Lineage trees for merged ILCs as in (**a**), represented with the population labels (left) and markers (right).

**Figure S6.**
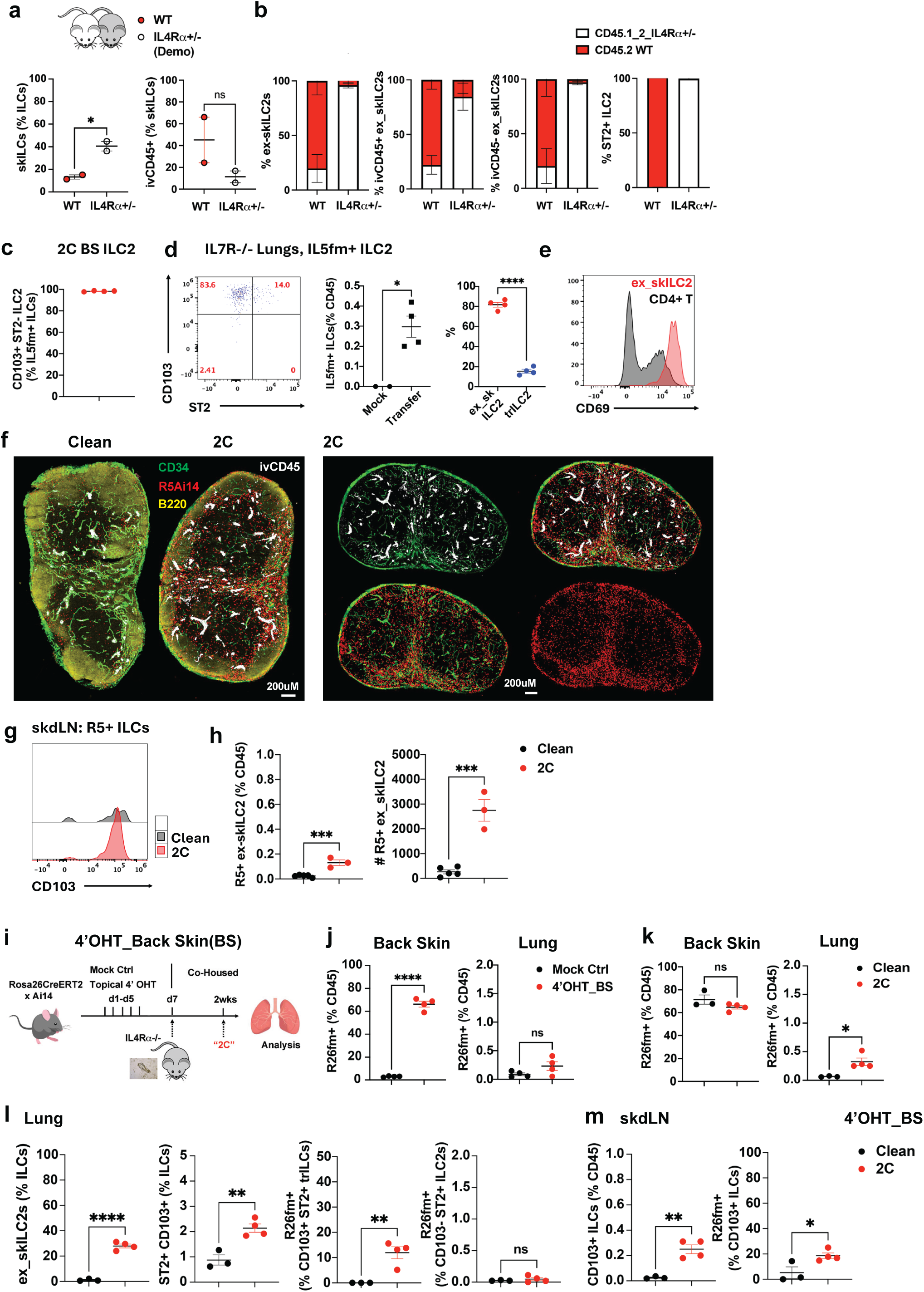
Tissue retention and adaptation of migratory skin-like ILCs. **a.** Parabiosis model of surgically linked 6-8wks old CD45.2 WT (red) and CD45.1/2 IL4Rα^+/–^ (Demo, white) mice. **b.** Graphs depict the percentage of ex-skILC2s and ivCD45^+^ ex-skILC2s in each mouse, and the ratio for the contribution of donor-derived cells for ivCD45^+^ ex-skILC2, ivCD45^-^ ex-skILC2 and ST2^+^ ILC2 in the lungs of each parabiont. **c.** Percentage of CD103^+^ ILC2s in 2C back skin. **d.** FACS plot (left) and quantification (right) of Il5^fm+^ ILC2s from skin that were transferred into IL7R^-/-^ host. Flow plot highlights the appearance of skILC2s in the lungs at 2wks. Percentage of IL5^fm+^ ex-skILC2s in lungs compared with lung-adapted ST2^+^CD103^+^ trILC2 (right). **e.** CD69 expression on transferred ex-skILC2s compared with CD4^+^ T cells. **f.** Confocal imaging for clean versus 2C skin-draining inguinal LNs (skdLN), cell distribution, and accumulation of lL5^fm+^ ex-skILC2s. **g.** Expression of CD103 by R5 ILCs in clean and 2C skdLN, **h.** Percentage and cell count of ex-skILC2s in skdLN. **i.** Experimental model for topical 4’OHT back skin painting (4’OHT_BS), followed by co-housing with Demodex mite donor. **j.** Labelling efficiency for skin (left) and lung (right) comparing mock and 4’OHT treated skin. **k.** pre-labelled (R26^fm+^) CD45^+^ skin cells from back skin and lungs in indicated groups. **l, m.** Percentage of ex-skILC2s, CD103^+^ ST2^+^ trILC2 and R26^fm+^ trILC2 and CD103^−^ ST2^+^ ILC2s of skin origin in clean and 2C lungs **(l)** and skdLN **(m)**. *p< 0.05, **p< 0.01, ***p<0.001, ****p<0.0001 by unpaired t-test. n=3-6 biological replicates/group, representative of 1-2 independent experiments.

**Figure S7.**
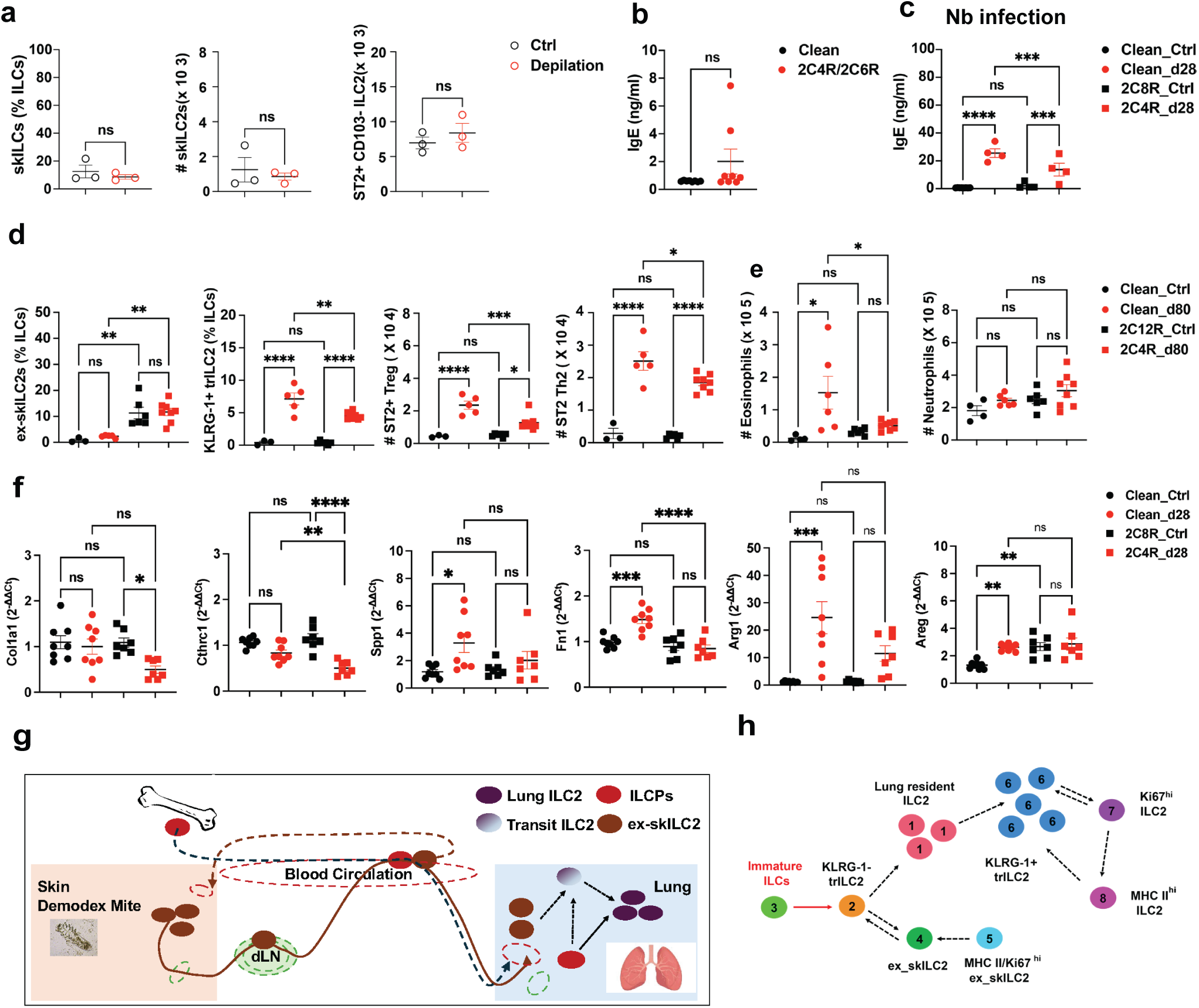
Prior Demodex experience lowers subsequent allergic inflammation in a heterologous parasitic challenge. **a.** Analysis of lung from shaved (control) vs depilated mice for percentage and cell count of skILC2s and ST2^+^ ILC2 in lungs 15 days after depilation. **b.** ELISA analysis for serum IgE from clean and 2C4R/2C6R mite-experienced mice. **c.** ELISA analysis for serum IgE for indicated groups from the *Nb* infection experiment. **d, e.** Percentage of ex_skILC2 and KLRG-1^+^ trILC2 and cell count for ST2^+^ Treg, ST2^+^ Th2, eosinophils and neutrophils from *Nb* infected lungs from the indicated group at d80 post infection. **f.** Gene expression by qPCR from lung tissue genes associated with fibroblast and tissue repair, gene list in **Table S2**. **g.** Conceptual figure of tissue egress and retention of migratory skin-like ILC2s in lungs. **h.** Summary figure of re-organized lung ILC2 landscape in response to helminth infection *p< 0.05, **p< 0.01, ***p<0.001, ****p<0.0001 by unpaired t-test (**a, b**) or one-way ANOVA (**c-f**). n=3-6 biological replicates/group, representative of 1-2 independent experiments.

---

**Table S1.**
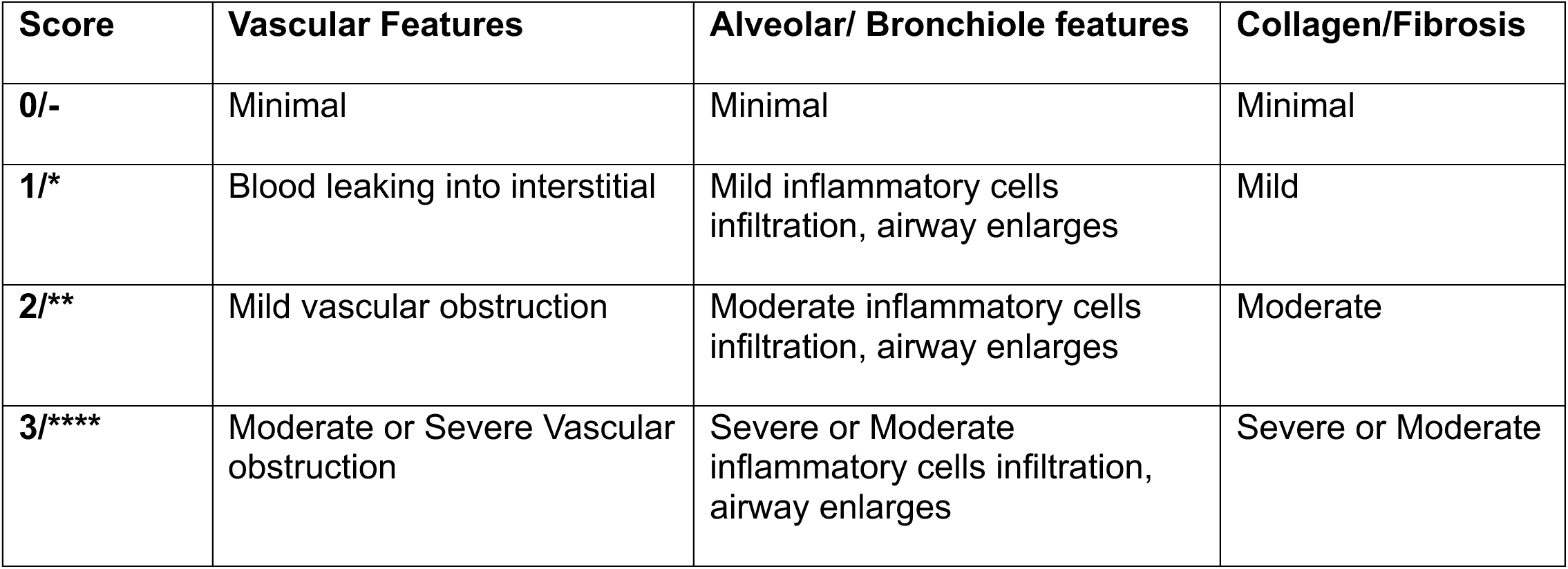
Lung pathology scoring system.

**Table S2.**
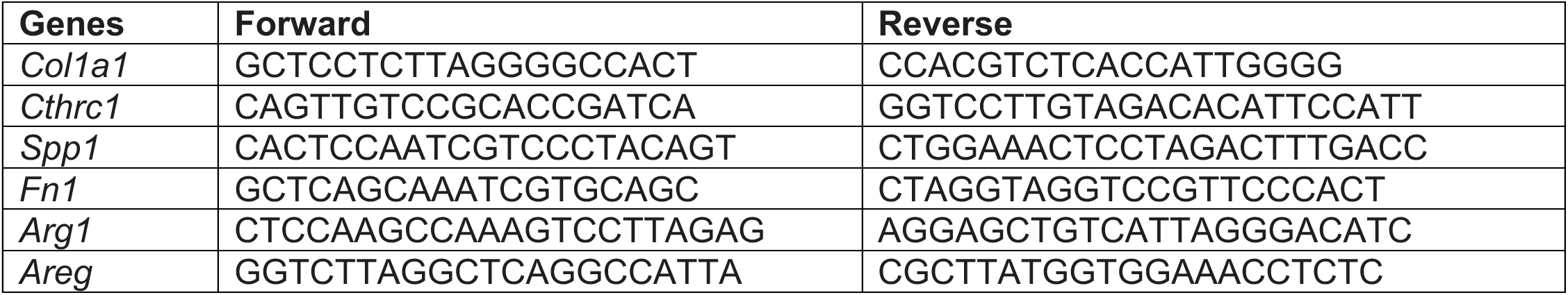
Real-time qPCR Primers: 5’ 3’.

**Table S3** Differential gene list of skin ILC2s

